# DNA sequence homology induces cytosine-to-thymine mutation by a heterochromatin-related pathway

**DOI:** 10.1101/115501

**Authors:** Eugene Gladyshev, Nancy Kleckner

## Abstract

In the genomes of many eukaryotes, including mammals, highly repetitive DNA is normally associated with histone H3 lysine-9 di-/trimethylation (H3K9me2/3) and C5-cytosine methylation (5mC) in the context of heterochromatin. In the fungus Neurospora crassa, H3K9me3 and 5mC are catalyzed, respectively, by a conserved SUV39 histone methylase DIM-5 and a DNMT1-like cytosine methylase DIM-2. Here we show that DIM-2 can also mediate cytosine-to-thymine mutation of repetitive DNA during the pre-meiotic process known as Repeat-Induced Point mutation (RIP) in N. crassa. We further show that DIM-2-dependent RIP requires DIM-5, HP1, and other heterochromatin factors, implying the involvement of a repeat-induced heterochromatin-related process. Our previous findings suggest that the mechanism of homologous repeat recognition for RIP involves direct pairwise interactions between co-aligned double-stranded (ds) DNA segments. Our current findings therefore raise the possibility that such pairing interactions may occur not only in pre-meiotic but also in vegetative cells, where they may direct heterochromatin assembly on repetitive DNA. In accord with this possibility, we find that, in vegetative cells of N. crassa, our model repeat array comprising only four 674-bp sequence copies can trigger a low level of DIM-5-dependent 5meC. We thus propose that homologous dsDNA/dsDNA interactions between a small number of repeat copies can nucleate a transient state of heterochromatin and that, on longer repeat arrays, such interactions lead to the formation of stable heterochromatin. Since the number of possible pairwise dsDNA/dsDNA interactions will scale non-linearly with the number of repeats, this mechanism provides an attractive way of creating the extended domains of constitutive heterochromatin found in pericentromeric and subtelomeric regions.

---

In the genomes of many eukaryotes, including mammals, highly repetitive DNA is normally associated with histone H3 lysine-9 di-/trimethylation (H3K9me2/3) and C5-cytosine methylation (5mC) in the context of heterochromatin^1-6^. In the fungus *Neurospora crassa*, H3K9me3 and 5mC are catalyzed, respectively, by a conserved SUV39 histone methylase DIM-5 and a DNMT1-like cytosine methylase DIM-2^7^. Here we show that DIM-2 can also mediate cytosine-to-thymine mutation of repetitive DNA during the pre-meiotic process known as Repeat-Induced Point mutation (RIP) in *N. crassa*^8–9^. We further show that DIM-2-dependent RIP requires DIM-5, HP1, and other heterochromatin factors, implying the involvement of a repeat-induced heterochromatin-related process. Our previous findings^10-12^ suggest that the mechanism of homologous repeat recognition for RIP involves direct pairwise interactions between co-aligned double-stranded (ds) DNA segments. Our current findings therefore raise the possibility that such pairing interactions may occur not only in pre-meiotic but also in vegetative cells, where they may direct heterochromatin assembly on repetitive DNA. In accord with this possibility, we find that, in vegetative cells of *N. crassa*, our model repeat array comprising only four 674-bp sequence copies can trigger a low level of DIM-5-dependent 5meC. We thus propose that homologous dsDNA/dsDNA interactions between a small number of repeat copies can nucleate a transient state of heterochromatin and that, on longer repeat arrays, such interactions lead to the formation of stable heterochromatin. Since the number of possible pairwise dsDNA/dsDNA interactions will scale nonlinearly with the number of repeats, this mechanism provides an attractive way of creating the extended domains of constitutive heterochromatin found in pericentromeric and subtelomeric regions.

The phenomenon of Repeat-Induced Point mutation (RIP) in *Neurospora crassa* occurs in the haploid germline nuclei that continue to divide by mitosis in preparation for karyogamy and ensuing meiosis^8,9^. During RIP, duplications of chromosomal DNA longer than a few hundred base-pairs are detected and subjected to strong C-to-T mutation over the extent of shared homology^7^. DNA duplications are recognized largely irrespective of their particular sequence, transcriptional capacity, or relative/absolute positions in the genome. In some filamentous fungi, the analogous process leads to cytosine methylation rather than mutation^13^. Previously, we showed that RIP did not involve the canonical homology-recognition pathway mediated by MEI-3, the only RecA homolog in *N*. crassa^10,12^. We further showed that RIP could sense the presence of weakly similar DNA sequences as long as those sequences shared a series of interspersed homologous base-pair triplets spaced at intervals of 11 or 12 base-pairs^10^. This pattern is consistent with a mechanism of recombination-independent homology recognition that involves interactions between co-aligned double-stranded DNA molecules^10,12^.

In our earlier work, we had developed a sensitive RIP tester construct comprising one endogenous and one closely-positioned ectopic copy of an arbitrarily-chosen 802-bp segment of the *N. crassa* genome^10^ (Fig. 1b; Supplementary Information Fig. 1a). This construct triggered strong RIP in a wildtype genetic background: specifically, the sample of 24 progeny spores was found to contain 3163 mutations in the endogenous (“left”) repeat copy, 3153 mutations - in the ectopic (“right”) repeat copy, and 524 mutations - in the endogenous 600- bp segment of the linker region (Fig. 1d: data replotted from ref. 10 as “*rid*+/+, *dim*-2+/+”). We now confirm that all of the above mutations, including those of the linker region, were induced by the presence of DNA homology: if the ectopic repeat copy is specifically omitted, no RIP activity is detected (Fig. 1c).

**Figure 1.**
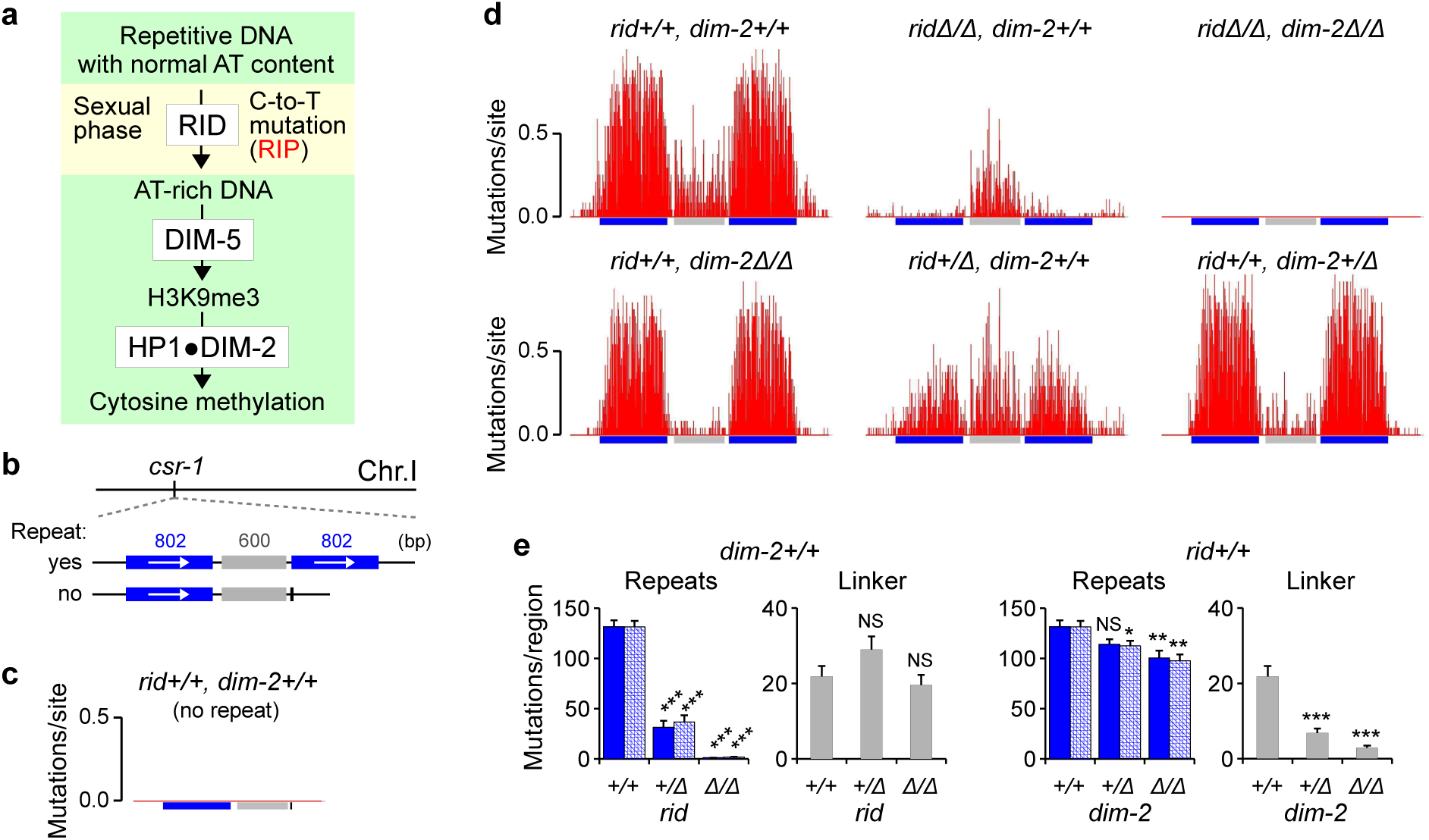
Cytosine methylase DIM-2 can mediate RIP. **a**, The canonical constitutive heterochromatin pathway in *Neurospora crassa*^7^. The pre-meiotic phenomenon of Repeat-Induced Point mutation (RIP) converts repetitive DNA with regular AT content into AT-rich DNA. AT-rich sequences recruit DIM-5 (the SUV39 histone methylase that catalyzes H3K9me3) in a complex with other proteins (most notably DDB1, CUL4, DIM-7 and DIM-9). Heterochromatin protein 1 (HP1) recognizes DIM-5-mediated H3K9me3 and directly recruits DIM-2 (the DNMT1-like C5-cytosine methylase) to DNA. **b**, The 802-bp tester construct contains one endogenous (“left”) and one closely-positioned ectopic (“right”) copy of a *Neurospora* genomic DNA segment (ref. 10). The “no-repeat” construct specifically lacks the ectopic repeat copy. Each construct is integrated by homologous recombination as the replacement of the *cyclosporin-resistant-1 (csr-1)* gene (Supplementary Information Fig. 1a). **c**, RIP mutation does not occur in the absence of the ectopic repeat copy. Cross X1[N=48] (the number of analyzed spores is provided in square brackets). **d**, RIP mutation profiles of the 802-bp construct in crosses with different combinations of *rid* and *dim-2* genotypes. Top row, from left to right: crosses X2[N=24], X3[N=48] and X4[N=48]; bottom row, from left to right: crosses X5[N=24], X6[N=24] and X7[N=24]. The number of mutations is reported per site per spore (corresponding to the probability of mutation). The wildtype cross (X2) was published previously (ref. 10). **e**, The mean (per-spore) number of RIP mutations for the crosses in **d**, measured in the three regions of interest: the 802-bp endogenous repeat copy (solid blue), the 802-bp ectopic repeat copy (hatched blue), and the 600-bp portion of the linker region, as defined in **b** (gray). Parental genotypes are provided in Supplementary Information Table 2. Error bars represent standard error (s.e.m.). Distributions of per-spore mutation counts are compared by the Kolmogorov-Smirnov (K-S) test (ref. 10). Each pair-wise K-S test uses the wildtype cross (X2) as a reference. *** P ≤ 0.001, ** 0.001 < P ≤ 0.01, * 0.01 < P ≤ 0.05, NS 0.05 < P.

In *N. crassa*, RIP has long been known to require a specialized putative C5-cytosine methylase RID (“RIP defective”)^14^. However, the genomes of several fungal species contain signatures of RIP-like mutation but do not encode any apparent homologs of RID, hinting at the possibility that RIP could be mediated by other factors^15,16^. In accord with this conjecture, we now find that our 802-bp tester construct can induce substantial mutation in the absence of RID (Fig. 1d,e: “*rid*Δ/Δ, *dim*-2+/+”). Intriguingly, while the expected (mean) number of mutations in the repeated regions is decreased by nearly two orders of magnitude in the *rid*Δ/Δ background, the intervening linker region is still mutated at essentially the wildtype level. Importantly, the findings above (Fig. 1c) imply that all of these RID-independent mutations are induced by DNA homology.

In addition to RID, *N. crassa* encodes another C5-cytosine methylase, DIM-2 (“Defective in methylation-2”)^17^, which catalyzes all known 5meC in this organism, as it occurs in several contexts, including newly-integrated multi-copy transgenes^18-20^. We now find that all RID-independent RIP mutation of the 802-bp construct requires DIM-2: when both RID and DIM-2 are absent, mutation can no longer be detected (Fig. 1d,e: “*rid*(Δ/Δ, *dim*-2Δ/Δ”). Furthermore, when DIM-2 is absent but RID is present, the pattern of effects is the reciprocal of that observed when DIM-2 is present but RID is absent (above): the number of mutations declines strongly (by the factor of 7.6) in the linker region and only moderately (by the factor of 1.3) in the repeated regions (Fig. 1d,e: “*rid*+/+, *dim*-2(Δ/Δ”). Interestingly, both *rid*Δ and *dim-2Δ* appear haploinsufficient: when present in combination with a corresponding wildtype allele, each gene deletion decreases the number of mutations in the corresponding affected region(s) by the factor of 3 or more (Fig. 1d,e: “*rid*+/Δ, *dim*-2+/+” and “rid+/+, *dim-2*+/Δ”).

The above findings show that: (i) RID and DIM-2 can each individually mediate RIP; (ii) RID and DIM-2 together account for all RIP; (iii) in the context of the 802-bp construct, RID-mediated mutation targets predominantly the repeated sequences (but can also occur in the single-copy linker region in between), whereas DIM-2-mediated mutation targets predominantly the single-copy linker region (but can also occur at a much lower level in the repeats); and (iv) the effects of RID and DIM-2 are additive. Taken together, these results suggest that the RIP processes mediated by RID and DIM-2 are functionally distinct and, to a first approximation, independent of one another.

To confirm the distinct and complementary nature of the RID- and DIM-2-mediated pathways of RIP mutation, we defined pair-wise correlations, on a per-spore basis, between the numbers of mutations occurring in different segments of the 802-bp tester construct (Supplementary Information Fig. 2). In situations where RID and DIM-2 activities are both strong (genetic backgrounds “*rid*+/+, *dim-2*+/+” and “*rid*+/Δ, *dim*-2+/+”), the total number of mutations in the left and in the right repeat copy of each individual spore clone are strongly correlated. That is, if one repeat copy exhibits a certain number of mutations, so does the other copy, on a per-spore basis. This pattern is expected if the two repeat copies are mutated by the same process. In contrast, the number of mutations in the linker region correlates less strongly with the number of mutations in either repeat copy, as expected if “linker” mutations and “repeat” mutations were mediated by two separate processes (i.e. mediated by DIM-2 and RID, respectively).

In vegetative cells of *N. crassa*, DIM-2 is recruited to DNA by the physical association with Heterochromatin protein 1 (HP1)^7^. HP1 recognizes H3K9me3 which, in turn, is established by the SUV39 histone methylase DIM-5 (Defective in methylation-5)^7^. The above observations raised the possibility that DIM-2-mediated mutation might also require these same factors. As one test of this possibility, we investigated the functional relevance of DIM-5 for RIP. Indeed, when DIM-5 and RID are both absent, no mutation can be detected, even if DIM-2 is still available at the wildtype level (Fig. 2a: “*dim*-5Δ/Δ”). Furthermore, DIM-2-mediated mutation can be readily restored in the presence of a single functional *dim*-5+ allele (Fig. 2a: “*dim*-5Δ/+”).

**Figure 2.**
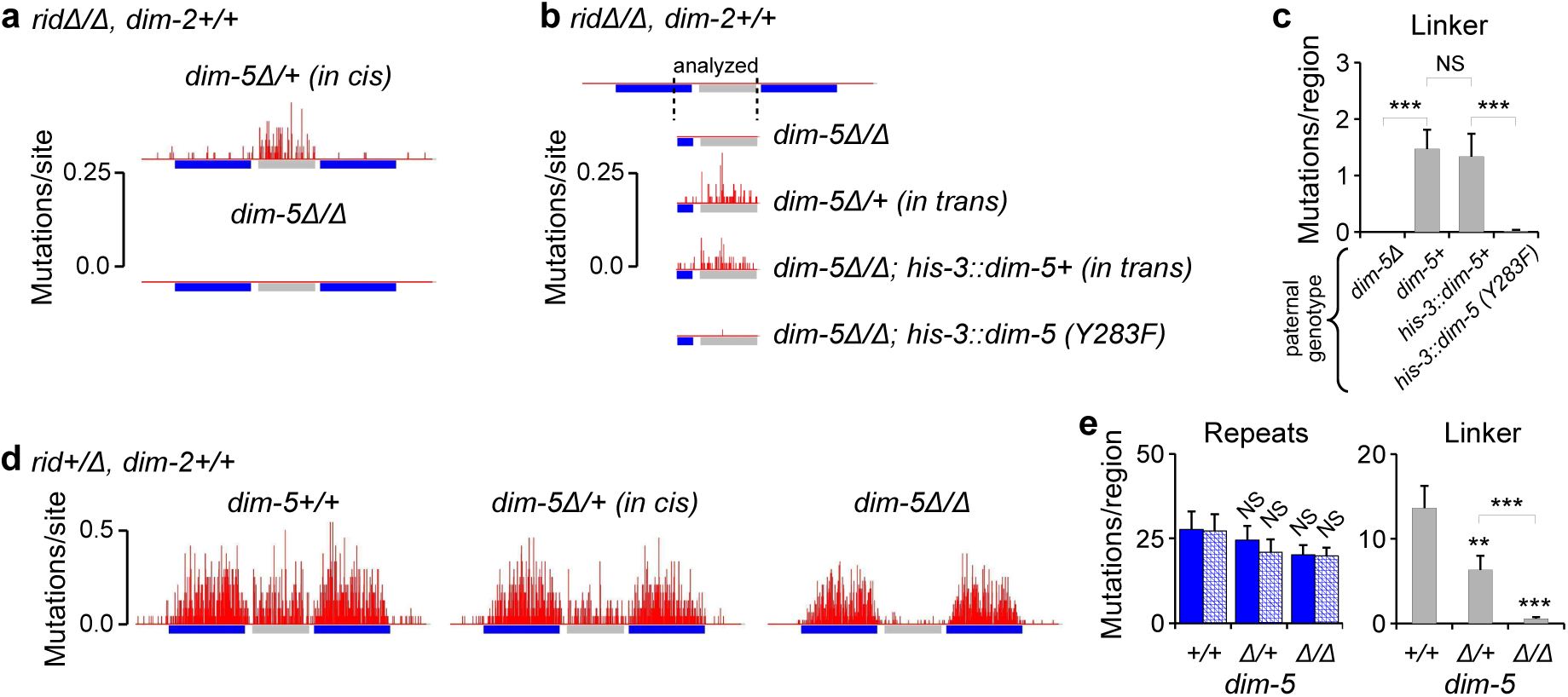
DIM-2-mediated RIP requires the SUV39 histone methylase DIM-5. Sexual development of female *dim*-5Δ strains is impaired in the presence of SET-7, an E(Z) H3K37 methylase encoded by the *set*-7 gene in *N. crassa*^22,23^. As RIP occurs normally in the absence of SET-7 (Supplementary Information Fig. 3b,c), the role of DIM-5 is assayed using *set*-7Δ female parents (Supplementary Information Table 2). **a**, RIP mutation profiles of the 802-bp construct. From top to bottom: crosses X8[N=60] and X9[N=60]. **b**, RIP mutation profiles of the 802-bp construct. From top to bottom: crosses X10[N=60], X11[N=60], X12[N=60] and X13[N=60]. Only the right-most portion of the endogenous repeat copy and the entire 600-bp segment of the linker region were analyzed by sequencing. **c**, The mean number of RIP mutations in the 600-bp segment of the linker region for crosses in **b** (analyzed as in Fig. 1d). **d**, RIP mutation profiles of the 802-bp construct. From left to right: crosses crosses X14[N=24], X15[N=24] and X16[N=48]. **e**, The mean number of RIP mutations for crosses in **d** (analyzed as in Fig. 1d). Error bars represent standard error (s.e.m.). Distributions of per-spore mutation counts are compared separately for each region of interest by the Kolmogorov-Smirnov test (as in Fig. 1d). *** P ≤ 0.001, ** 0.001 < P ≤ 0.01, * 0.01 < P ≤ 0.05, NS 0.05 < P. Parental genotypes are provided in Supplementary Information Table 2.

Additional experiments confirm and extend this conclusion. In the heterozygous *dim*-5Δ/+ cross analyzed above (Fig. 2a), the male parent provided the wildtype *dim*-5+ allele together with the 802-bp construct, whereas the female parent provided the null *dim*-5Δ allele. In such a configuration (where the functional *dim*- 5+ allele and the tester construct were supplied *“in cis”*; Supplementary Information Fig. 1c), DIM-5 could, in principle, have acted on the repeat construct in vegetative cells, prior to fertilization and the onset of the pre-meiotic stage during which RIP occurs. To investigate if the role of DIM-5 in RIP could be fulfilled after fertilization, when nuclei of both parental strains are brought into a common cytoplasm, we assayed the RIP-promoting activity of DIM-5 in a cross between a repeat-carrying maternal *dim*-5Δ strain and a repeat-less paternal strain that supplied a functional *dim*-5+ allele (i.e., the *dim*-5+ allele and the tester construct were provided “*in trans*”). We find that “*cis*” and “*trans*” configurations result in the comparable levels of RIP, regardless of whether the paternal *dim*-5+ allele is present in the endogenous or in the ectopic locus (Fig. 2b,c). These results imply that DIM-5 was acting after fertilization. Furthermore, nearly all mutation observed in the “*trans*” configuration can be abolished by a single amino-acid change in DIM-5 that eliminates a catalytically important tyrosine residue^21^ (Fig. 2b,c: “*dim*-5Δ/Δ; his-3::*dim*-5 (Y283F)”). The fact that the mutant DIM-5(Y283F) protein still exhibits some very weak RIP-promoting activity *in vivo* corresponds to the fact that it also retains weak residual catalytic activity *in vitro*^21^. Thus, DIM-5 appears to exert its role in RIP by its canonical H3K9 trimethylase activity.

We also examined the effect of decreasing DIM-5 levels in the presence of RID (Fig. 2d,e). The complete lack of DIM-5 confers a strong and specific phenotype in the linker region (a 24.3-fold decline) while having a much more modest impact on the repeated sequences (a 1.4-fold decline, similar to the 1.3-fold decline observed in repeats in the absence of DIM-2; Fig. 1d,e). These findings provide further evidence that RID-mediated RIP is largely independent of DIM-5.

The finding that RIP could be mediated by the DIM-5/DIM-2 pathway raised the possibility that *Neurospora* HP1 (encoded by the *hpo* gene) might also be involved in RIP. In an attempt to test this idea, we found that the lack of HP1 precluded meiotic spore formation, even in the *set*-7Δ background that normally permitted sexual development in the absence of DIM-5 ^22,23^. This phenotype likely reflects a broader requirement for HP1 during the sexual phase in *N. crassa*. However, we were able to find a combination of two *hpo*Δ strains that produced some limited amount of late-arising spores. One of these strains (C135.3, used as a female parent) lacked the type II topoisomerase-like protein SPO11 that normally catalyzes the formation of double-strand DNA breaks in meiosis yet is fully dispensable for RIP^10,12^. Our analysis shows that the apparent level of RIP mutation is strongly reduced in the *hpo*Δ/Δ background (Supplementary Information Fig. 4c). However, (i) substantial RIP still occurs in the complete absence of HP1 (Supplementary Information Fig. 4b), and (ii) the relative levels of RIP in the linker region are very similar between the *hpo*Δ/Δ and *dim-2*Δ/Δ backgrounds (Supplementary Information Fig. 4f). Thus, we infer that HP1 is also involved in the DIM-5/DIM-2 pathway for RIP.

We next investigated the roles of two additional components of the *Neurospora* heterochromatin pathway: CUL4 (Cullin 4) and DDB1 (Damage specific DNA binding protein 1, also known as DNA damage-binding protein 1), which together comprise the core of the Cullin4-RING finger ligase CRL4^7^. CUL4 and DDB1 are both required for all DIM-5-mediated H3K9me3 in *Neurospora* vegetative cells^7^. We have found that the lack of either protein abrogates the normal development of perithecia (even when examined in various *set*-7Δ, *spoil* backgrounds), thus precluding our standard analysis of RIP. However, instead, we were able to assay the role of CUL4 and DDB1 in the two heterozygous conditionns (*cul*4+/Δ and *ddbi*+/Δ; Supplementary Information Fig. 4d-f). Our results show that, while the overall levels of RIP are substantially decreased in both situations, the pattern of changes is diagnostic of the specific defect in the DIM-5/DIM-2-mediated process, i.e. a stronger reduction of mutation in the linker region as compared to a more moderate reduction in the repeats (Supplementary Information Fig. 4d-f). Taken together, the above results suggest that RIP can be mediated by a process that does not require RID, and in which DIM-5, HP1, DIM-2 and CUL4/DDB1 act in a single heterochromatin-related pathway.

The next issue raised by these findings concerned the homology requirements for the newly discovered heterochromatin-related pathway of RIP. Our published results^10^ indicated that the absence of MEI-3 (the only RecA protein in *N. crassa*) did not detectably affect the patterns of mutation in a tester construct comprising two direct repeats of 800 base-pairs and separated by a single-copy region of 206 base-pairs. Thus, previous experiments provided no indication that the canonical recombination-mediated mechanism of homology recognition (mediated by MEI-3) played a role in either the RID- or the DIM-2-mediated pathways. We have also previously shown^10^ that in the wildtype situation, when both pathways are active, RIP can detect the presence of weak interspersed homology, as long as that homology obeyed certain rules, e.g, comprised short islands of homology (≥3 base-pairs) spaced at regular intervals of 11 (or 12) base-pairs along a pair of participating chromosomal DNA segments (Fig. 3b). This and other findings led us to a model in which homology recognition for RIP involved direct interactions between co-aligned DNA duplexes^10^. One series of experiments that led to this conclusion utilized repeat constructs in which 200-bp regions of interspersed homology (e.g., formed between light blue/magenta segments in Fig. 3a,b) were adjacent to a 220-bp region of perfect homology (formed between two blue segments in Fig. 3a,b)^10^. These direct “200+220” repeats were linked by a single-copy region of 537 base-pairs. The 220-bp region of perfect homology was incorporated to provide a stable, permanent point of interaction, thereby facilitating the detection of weak effects induced by interspersed homologies. In this context, different homology patterns could be created by manipulating only the “right” 200-bp segment while leaving the rest of the construct unchanged.

**Figure 3.**
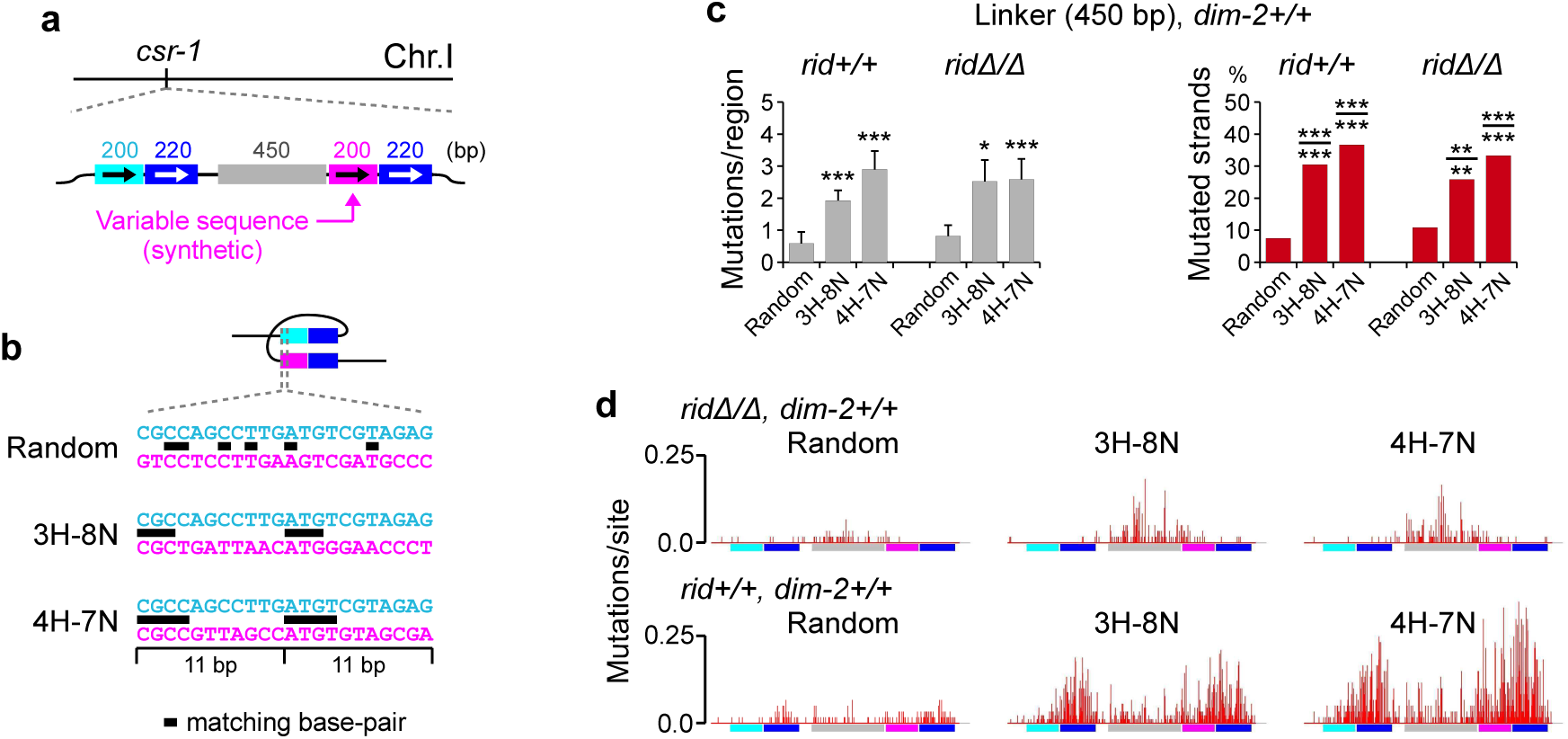
DIM-2-mediated RIP responds to the presence of weak interspersed homology. **a**, Recognition of interspersed homologies by RIP is assayed using previously published repeat construct (ref. 10), in which 220 base-pairs of perfect homology (between two blue regions) are adjacent to 200 base-pairs of interspersed homology (between light-blue and magenta regions). Different patterns of interspersed homology are created by varying the nucleotide sequence of the magenta region. **b**, Patterns of weak interspersed homology contain short homologous units of three (“3H-8N”) and four (“4H-7N”) base-pairs spaced at regular intervals of 11 base-pairs. “Random” homology corresponds to a 200-bp fragment of the GFP coding sequence (ref. 10). **c**, RIP mutation in the 450-bp segment of the linker region analyzed for conditions in **d**. RIP is quantified as the mean number of mutations (as in Fig. 1d; left panel) and as the percentage of mutated DNA strands (as in ref. 10, right panel). The following numbers of mutated/wildtype DNA strands were obtained: 9/111, 55/125, 44/76 (*rid*+/+, from left to right) and 13/107, 31/89, 40/80 (*rid*Δ/Δ, from left to right). Pair-wise differences in percentages of mutated strands are tested for significance as raw numbers using the chi-squared homogeneity test (*P*-value is indicated above the line) and the Fisher's exact test (*P*-value is indicated below the line). Each instance of interspersed homology is compared to random homology in a corresponding genetic background. **d**, RIP mutation profiles for constructs with random versus weak interspersed homology. Top row, from left to right: crosses X17[N=60], X18[N=60] and X19 [N=60]; bottom row, from left to right (as reported previously in ref. 10): crosses X20[N=60], X21[N=90] and X22[N=60]. Error bars represent standard error (s.e.m.). Distributions of per-spore mutation counts are compared by the Kolmogorov-Smirnov test (as in Fig. 1d). *** P ≤ 0.001, ** 0.001 < P ≤ 0.01, * 0.01 < P ≤ 0.05, NS 0.05 < P. Parental genotypes are provided in Supplementary Information Table 2.

We now used this same repeat system to address specifically the homology requirements for DIM2-mediated RIP. We first re-analyzed our published data, obtained in the *rid*+/+, *dim*-2+/+ background (ref. 10), with respect to mutation of the 450-bp segment of the linker region (Fig. 3a), which was expected to differentially report the effects of the DIM-5/DIM-2-mediated process. The analyzed instances of interspersed homology involve homologous units of 3 or 4 base-pairs spaced at regular intervals of 11 base-pairs (Fig. 3b, patterns “3H-8N” and “4H-7N”, respectively). The presence of these homology patterns produces a significant increase in mutation throughout the 450-bp segment, suggesting that DIM-2-mediated RIP can also respond to weak interspersed homologies (Fig. 3c,d: *rid*+/+, *dim*-2+/+). We then asked whether these same homology patterns could promote RIP in the absence of RID, when only the DIM-5/DIM-2 pathway remains functional. Here we find again that RIP in the linker segment is increased significantly in the presence of the assayed interspersed homologies (Fig. 3c,d: *rid*Δ/Δ, *dim*-2+/+). Moreover, the levels of mutation in the linker segment induced in the presence and in the absence of RID by any given homology pattern appear similar, suggesting that the majority of the previously reported “linker” mutations^10^ were mediated by DIM-2. Taken together these findings suggest that the heterochromatin-related pathway of RIP can also be activated by weak interspersed homology.

In all repeat constructs analyzed thus far, DIM-5/DIM-2-mediated RIP occurred predominantly in the singlecopy linker region, between the repeated sequences. This pattern of mutation suggests that the presence of relatively short repeats can direct the activity of the heterochromatin-related process to nearby non-repetitive regions. To further explore this idea in a different context, we analyzed a construct containing four copies of the dsRed coding sequence (the “4x” array) inserted near the *his-3* gene (Fig. 4a). As a baseline, we first examined mutation of the 4x array by RID-mediated RIP, in the absence of DIM-2. We find that the repeat copies are strongly mutated, yet no spreading of mutations into the adjoining single-copy regions has occurred (Fig. 4b: “*rid*+/Δ, *dim*-2Δ/Δ”). We then examined mutation of the 4x array by the DIM-5/DIM-2 pathway, in the absence of RID. In a dramatic contrast, DIM-5/DIM-2-dependent mutations occur at comparable levels in the repeated and the adjoining non-repetitive regions (Fig. 4b: “*rid*Δ/Δ, *dim*-2+/+”). Taken together, these results suggest that (i) tandem repeat arrays represent a very effective trigger of RIP; and (ii) the DIM-5/DIM-2 pathway targets not only the repeated sequences (which are nevertheless essential for activating this pathway), but also the adjoining non-repetitive regions.

**Figure 4.**
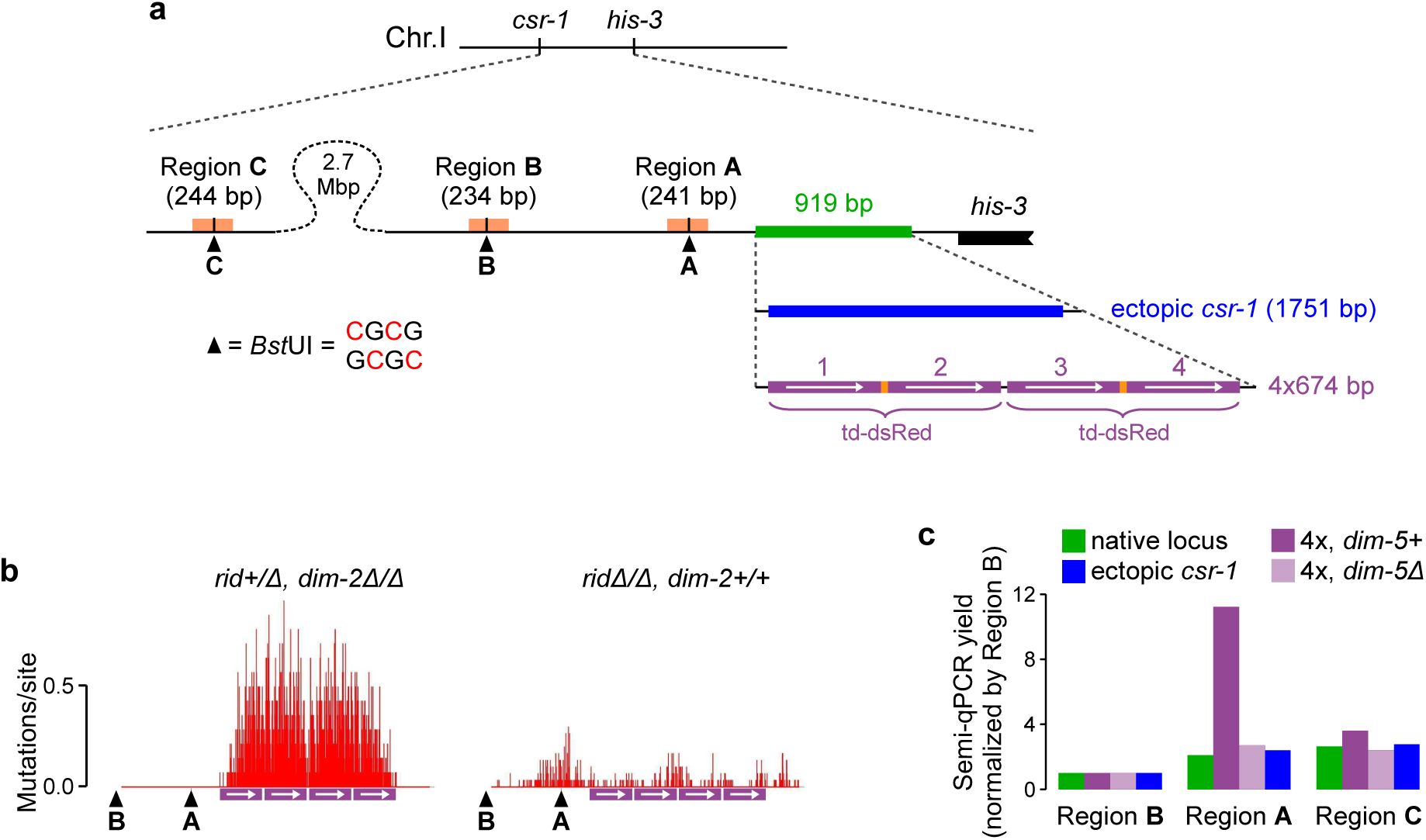
Recognition of short tandem repeat arrays by the DIM-5/DIM-2 pathway. **a**, Four copies of the dsRed coding sequence (the “4x” array, provided on the plasmid pEAG236A) are inserted by homologous recombination as the replacement of an endogenous genomic segment (green) near the *his-3* gene. Integration of the ectopic *csr-1* copy (analyzed in Fig. 5) is done analogously. All DNA segments are drawn to the same scale. The restriction site 5'-CGCG-3' can only be cut by *Bst*UI if none of its four cytosines (two cytosines on the top strand and two cytosines on the bottom strand) is methylated (Methods). The *Bst*UI site of Region C is located near the *csr-1* gene. **b**, RIP mutation profiles for the 4x array. Left: cross X26[N=14]; right: cross X27[N=30]. The mean number of mutations in the entire sequenced region is 174.93± 30.84 (cross X26) and 15.20±1.69 (cross X27). Positions of the two *Bst*UI restriction sites (“A” and “B”, as shown in **a**) are indicated. **c**, The assay detects the failure of cleavage of genomic DNA by *Bst*UI at the three specific sites (in Regions A, B and C, as shown in **a**). PCR yields for Regions A and C are normalized by the corresponding PCR yield for Region B. Unnormalized PCR yields are presented in Supplementary Information Fig. 6b. Assayed strains: FGSC#9720 (native locus), T485.4h (4x, *dim*-5+), T486.3h (4x, *dim*-5Δ) and T402.1h (ectopic *csr-1).* The analyzed repeat constructs have never been subjected to RIP, having been introduced by transformation and then propagated only by the vegetative growth (Supplementary Information Table 1; Methods).

All of the above analyses examined DIM-2-mediated RIP as triggered by closely-positioned repeats. It was of interest to determine whether this pathway could also target repeats at widely-separated genomic positions. To address this question, we designed a sensitive genetic system that can detect RIP between a single pair of homologous sequences located 2.7 Mbp apart on the same chromosome (Fig. 5a; also ref. 12). The *csr-1* gene encodes a cyclophilin protein that has a high affinity for cyclosporin A, and the presence of a single active copy of *csr-1* is sufficient to render a *Neurospora* strain cyclosporin-sensitive. Thus, if two *csr-1* copies are present, one of which is active and the other is not, RIP mutation of the active copy (induced by the ectopic inactive copy) will generate cross progeny resistant to cyclosporin (Fig. 5b). We find that in the absence of the ectopic copy, no cyclosporin-resistant progeny could be detected among nearly 5×10^5^ spores produced by a cross between two wildtype strains, implying that the frequency of spontaneous mutation of the *csr-1* gene is extremely low. In contrast, in the presence of the ectopic copy, a large proportion of cross progeny becomes cyclosporin-resistant (Fig. 5c). This proportion decreases by nearly four orders of magnitude in the absence of both RID and DIM-2. Rare cyclosporin-resistant progeny still emerge in the *rid*Δ/Δ, *dim*-2Δ/Δ condition. However, as these spore clones all carry the same G-to-A mutation that is already present in the parental ectopic copy, these rare events can likely be attributed to gene conversion between the endogenous and the ectopic copies (Fig. 5c,d). Finally, when DIM-2 is present and RID is absent, cyclosporin-resistant progeny are produced at a level that is one order of magnitude higher than the gene-conversion background. Sequence analysis of null *csr-1* alleles reveals the occurrence of only C-to-T and G-to-A mutations, confirming that these alleles were indeed produced by RIP (Fig. 5d). Similarly to the pattern of mutations observed at closely-positioned repeats (above), RID-mediated mutations at widely-separated repeats remain confined to the duplicated region, whereas DIM-2-mediated mutations tend to spread into the adjoining single-copy regions (Fig. 5d, Supplementary Information Fig. 5). An additional interesting difference between the two mutational pathways is the even stronger preference for CpA sites associated with DIM-2-mediated RIP (Fig. 5e). Taken together, these results demonstrate the capacity of the heterochromatin-related pathway to mediate mutation of DNA repeats that are separated by large genomic distances.

**Figure 5.**
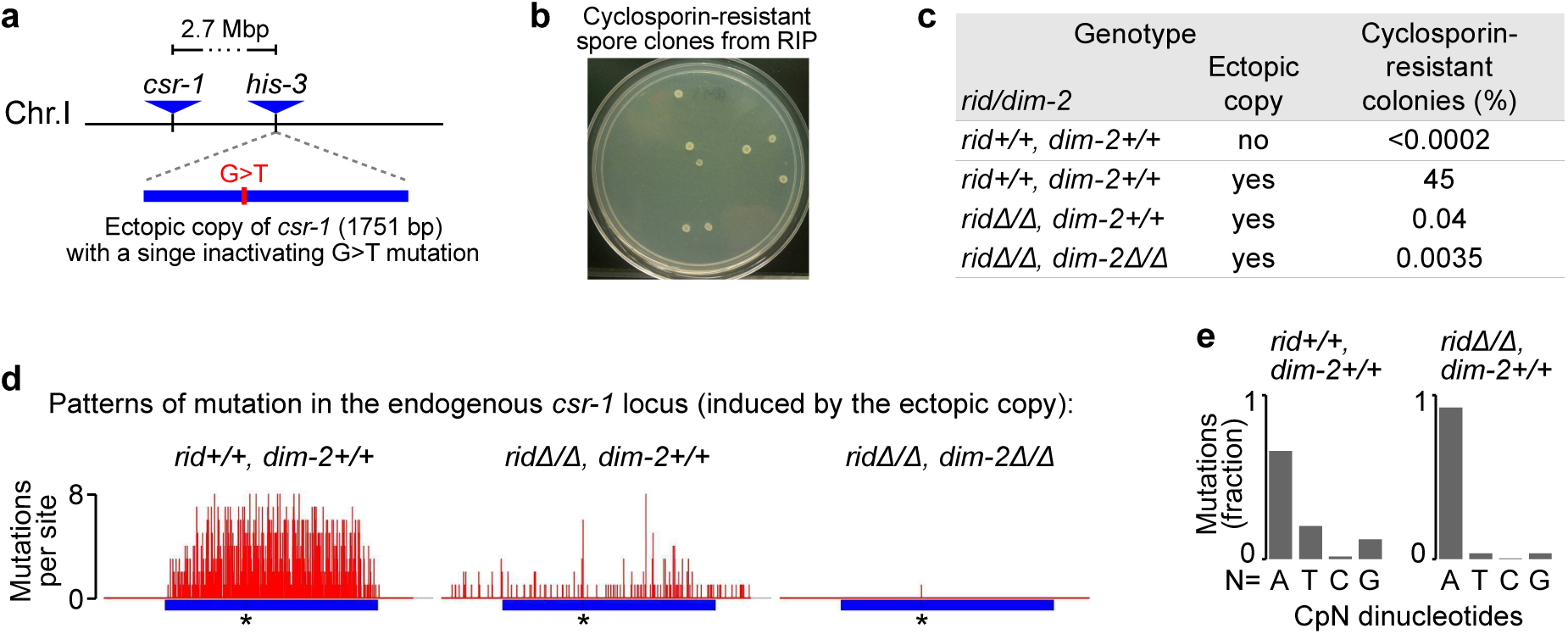
DIM-2-mediated RIP can target repeats at widely separated genomic positions. **a**, A pair of widely-separated repeats of the *csr-1* gene: the inactive ectopic copy of *csr-1* is inserted near the *his-3* gene, 2.7 million base-pairs away from the endogenous *csr-1* locus. **b**, Inactivation of the endogenous *csr-1* gene by RIP produces cyclosporin-resistant spore clones. **c**, The percentage of cyclosporin-resistant progeny produced in different genetic backgrounds. Because the ectopic repeat is present in only one of the two parental strains, the maximum expected level of cyclosporin-resistant progeny is 50 per cent. **d**, Patterns of RIP mutation observed in the endogenous *csr-1* locus. The per-site number of mutations is reported for sets of distinct *csr-1* alleles. “*” denotes the position of the inactivating G-to-A mutation in the ectopic copy. The number of replica crosses, the total number of sequenced cyclosporin-resistant progeny (N), and the number of distinct *csr-1* alleles are as follows: X23, *rid+/+, dim-2+/+* [1 cross, N=10 sequenced spores, 10 distinct alleles]; X24, *rid*Δ/Δ, *dim*-2+/+ [9 crosses, N=70 sequences spores, 34 distinct alleles]; X25, *rid*Δ/Δ, *dim*-2Δ/Δ [3 crosses, N=28 sequenced spores, 1 distinct allele]. We note that 20 cyclosporin-resistant progeny in the *rid*Δ/Δ, *dim*-2Δ/Δ background were likely produced by a single “jackpot” event that could have occurred early in one pre-meiotic lineage. **e**, Dinucleotide context of RIP mutations.

Overall, the observations presented above demonstrate that, in pre-meiotic cells of *Neurospora*, relatively short repeats can be recognized for RIP mutation by a heterochromatin-related pathway. Interestingly, this pathway targets not only the repeats themselves but also adjacent single-copy regions, another characteristic feature of heterochromatin assembly in vegetative/somatic cells^1-6^. Previous studies^10,11^, extended above, suggest that repeat recognition for this heterochromatin-related RIP pathway is triggered by pairwise dsDNA/dsDNA interactions. Taken together, these findings raise the possibility that dsDNA/dsDNA interactions might also be involved in the formation of heterochromatin in vegetative cells, but result in cytosine methylation instead of mutation. We investigated this possibility directly by analyzing our short repeat constructs for their ability to set up heterochromatin-related events in vegetatively-growing *Neurospora* cells.

The nature and requirements for heterochromatin formation on repetitive DNA in *N. crassa* have been extensively characterized^18-20^.^24^. In these studies, heterochromatin was detected only on long repeat arrays. For example, 25 tandem copies of the *al-1* gene were found to induce stable heterochromatin^20^. yet no signs of heterochromatin could be detected over an array containing four tandem copies of the *am* gene^24^. The same threshold effect can also be seen in other organisms, including mammals. For example, in the well-studied case of FacioScapuloHumeral Muscular Dystrophy Type 1 (FSHD1), the disease state is triggered by the contraction of the D4Z4 macrosatellite array from 100-11 to 10-1 copies, and this contraction is accompanied by the loss of heterochromatic silencing (as defined by the progressive removal of H3K9me3 and 5meC) as well as the concomitant reactivation of the D4Z4-embedded *DUX4* gene^25^.

The presence of an apparent repeat-number threshold raises an interesting possibility: heterochromatin may normally be nucleated on short repeat arrays but is then removed unless it can be stabilized by the presence of a larger number of repeats. If this process involves transient formation of heterochromatin, the diagnostic signal will be present in only a small fraction of cells and thus could not be easily detected by the bulk assays, e.g. analysis of H3K9me3 by chromatin immunoprecipitation (ChIP) or of 5meC by Southern blots or bisulfite conversion. In contrast, our short repeat constructs used for the analysis of RIP, if combined with a sensitive detection assay, could provide a model system for identifying such transient nucleation events.

We therefore chose to analyze the 4x array (Fig. 4a), which provides a suitable model of a short repeat array. To detect transient (rare) heterochromatin nucleation events, we utilized a sensitive PCR-based assay for C5-cytosine methylation, which is a diagnostic mark of heterochromatin in *N. crass^7^*. Specifically, we analyzed the state of a particular restriction site 5'-CGCG-3' that can be cleaved by *Bst*UI only if none of its four cytosines is methylated (Methods). This *Bst*UI site is located just to the “left” side of the integration position of the 4x array, in a single-copy genomic region that shows strong DIM-5/DIM-2 dependent RIP (Fig. 4a,b: “A”). We analyzed this site for the 5meC-mediated inhibition of cleavage under several conditions, using, as normalization controls, two additional *Bst*UI sites that were not expected to undergo heterochromatinization (Fig. 4a-c: “ B” and “C”). We find that *Bst*UI cleavage at the reporter site is prominently and specifically inhibited when the 4x array is present as indicated by enrichment of a diagnostic PCR product that spans the critical site (Fig. 4c: PCR Region A; unnormalized data are provided in Supplementary Information Fig. 6). Furthermore, there is no detectable evidence of repeat-dependent methylation when DIM-5 is absent, implying that the detected 5meC is dependent on this key heterochromatin factor. Detectable 5meC is absent in a control construct comprising a copy of the *csr-1* gene that is integrated instead of the 4x array (Fig. 4a). Importantly, the level of the diagnostic PCR signal induced by the 4x array in the presence of DIM-5 is low and corresponds to the 5meC frequency of *~*2-3*%*. We conclude that our assay is detecting rare (and thus likely transient) events of the repeat-induced heterochromatin formation in vegetative cells of *N. crassa*.

Our findings suggest that transient heterochromatinization can also occur on a short repeat array with only four relatively short repeat copies. These results have two important general implications. First, they support the idea that transient local events involving small numbers of repeat copies may underlie the formation of stable heterochromatin on long repeat arrays. Second, they suggest that the rules for repeat recognition during RIP, e.g. pairwise interactions between co-aligned dsDNA segments, could, analogously, underlie repeat recognition during heterochromatin assembly in vegetative/somatic cells.

A further question arises as to why the 4x array elicits a robust RIP mutation signal in pre-meiotic cells but only a weak signal, diagnostic of unstable heterochromatin, in vegetative cells. This difference likely reflects the combined effects of two factors. First, during RIP, a transient pairwise interaction involving a small number of repeat copies produces a permanent record of each event in the form of mutation. Second, each nucleus may undergo as many as 6-7 rounds of RIP over the course of several days. As a result, mutations resulting from multiple transient pairing and heterochromatinization events will be present in the accumulated form in the progeny spores that are finally analyzed.

The apparent connection between the repetitive nature of chromatin and its heterochromatic status was noted by Guido Pontecorvo in 1944 (ref. 26) and elaborated further by Douglas Dorer and Steven Henikoff^27^, with the suggestion that the presence of repeats might be recognized *per se*. However, current models of heterochromatin formation invoke repeat recognition via cis-acting proteins that recognize specific sequence motifs or small RNAs that associate with nascent, newly-transcribed RNA at the repetitive locus^1,3,6^. The presented findings provide evidence for a different molecular mechanism in which repeat recognition for heterochromatin formation involves direct, pairwise, dsDNA/dsDNA interactions between repeats.

We propose that heterochromatin formation in vegetative cells is nucleated at short repeat sequences by pairwise dsDNA/dsDNA interactions involving a small number of repeat units (Fig. 6). Stable heterochromatin on long repeat arrays could then arise via multiple nucleation events of this type as well as by extension of pairwise interactions outward from nucleation site(s). Given the pairwise nature of repeat recognition, the number of potential nucleation events will scale more-or-less exponentially with the number of repeat copies, thus giving an effective way of nucleating and extending heterochromatin over very large repeat arrays.

**Figure 6.**
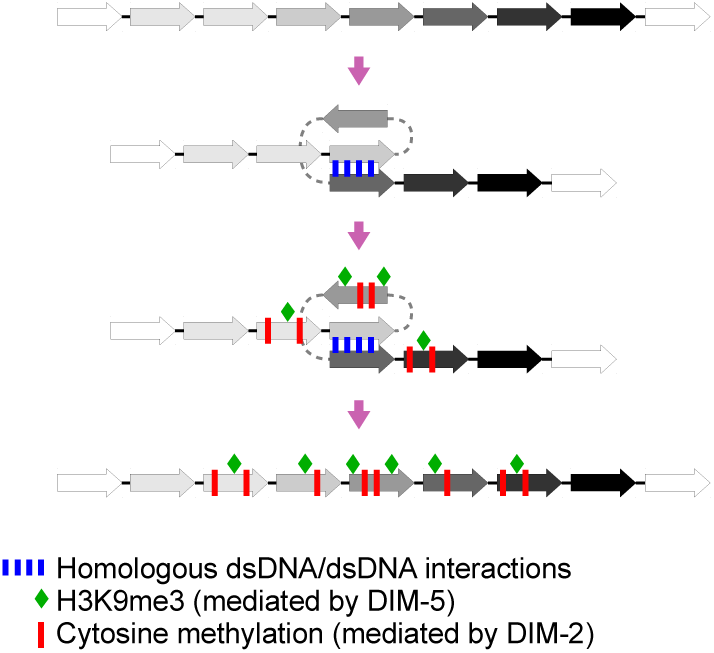
A general model for nucleation of heterochromatin by pairwise dsDNA/dsDNA interactions. In vegetative/somatic cells, a direct homologous dsDNA/dsDNA interaction between two co-aligned copies of a repeated sequence nucleates local assembly of heterochromatin as marked by H3K9me3 (green diamonds) and 5meC (red bars). These local assemblies are intrinsically unstable. However, on longer repeat arrays, multiple concomitant nucleation events may be stabilized to yield constitutive heterochromatin. In pre-meiotic cells of *N. crassa*, the same transient local interactions result in permanent DNA marks in the form of C-to-T mutations, instead of 5meC. Such mutations accumulate over several cell cycles to give a high level of RIP in analyzed spores.

The discovery by Michael Freitag, Eric Selker and colleagues that RIP in *N. crassa* required RID^14^ created a conundrum: RIP involves cytosine-to-thymine mutation, whereas RID has all the conserved features of a C5-cytosine methylase^12,14^. Thus, it was proposed that RIP mutation might occur as a two-step process, in which cytosine methylation (by RID) could be followed by 5meC deamination (by another enzymatic activity)^14^.

This idea was supported by the fact that in another filamentous fungus, *Ascobolus immersus*, pre-meiotic recognition of repeats leads to cytosine methylation by Masc 1 (the only RID homolog in *A. immersus*) without mutation^13^. An alternative proposal was also put forward by which RID alone could mediate C-to-T mutation, via modulation of its catalytic activity^14^. Our current findings reveal that RIP can be mediated by the canonical cytosine methylase DIM-2. In vegetative cells of *N. crassa*, DIM-2 catalyzes all known cytosine methylation, without obvious deamination. Interestingly, DIM-2 methylates cytosines in all dinucleotide contexts^28^, whereas DIM-2-mediated mutation exhibits a very strong preference for CpA dinucleotides (Fig. 5e), pointing to a distinct substrate specificity for deamination as compared to methylation. These findings are fully compatible with a two-step mechanism for RIP. Overall, by this hypothesis, the presence of DNA homology would trigger cytosine methylation (by either DIM-2 or RID), and resulting 5meC would then be converted to thymines by a deaminase that becomes active specifically during the pre-meiotic phase. It can be noted, however, that in vegetative cells, DIM-2 can also mediate methylation of non-repetitive coding parts of the genome^29^. If that same process takes place in pre-meiotic nuclei, then a two-step mechanism would further require the deamination step to be dependent on the presence of homology as well. A one-step mechanism involving pre-meiotic modulation of the catalytic activity of DIM-2 would not have such a requirement.

Finally, we note that the notable spread of DIM-5/DIM-2-mediated epigenetic and genetic modifications from repetitive DNA into neighboring single-copy regions implicates this homology-directed pathway in the heterochromatic silencing and the accelerated evolution of pathogenic genes that are frequently associated in clusters with repetitive elements in the genomes of many filamentous fungi^16,30^.

## ACKNOWLEDGEMENTS

We thank Denise Zickler for critically reading the manuscript. This work was supported by the grant GM044794 from the National Institutes of Health to N.K. and research fellowships from The Helen Hay Whitney Foundation, The Howard Hughes Medical Institute and Charles A. King Trust to E.G.

## AUTHOR INFORMATION

The authors declare no competing financial interests.

## SUPPLEMENTARY INFORMATION FIGURE LEGENDS

**Supplementary Information Figure 1.**
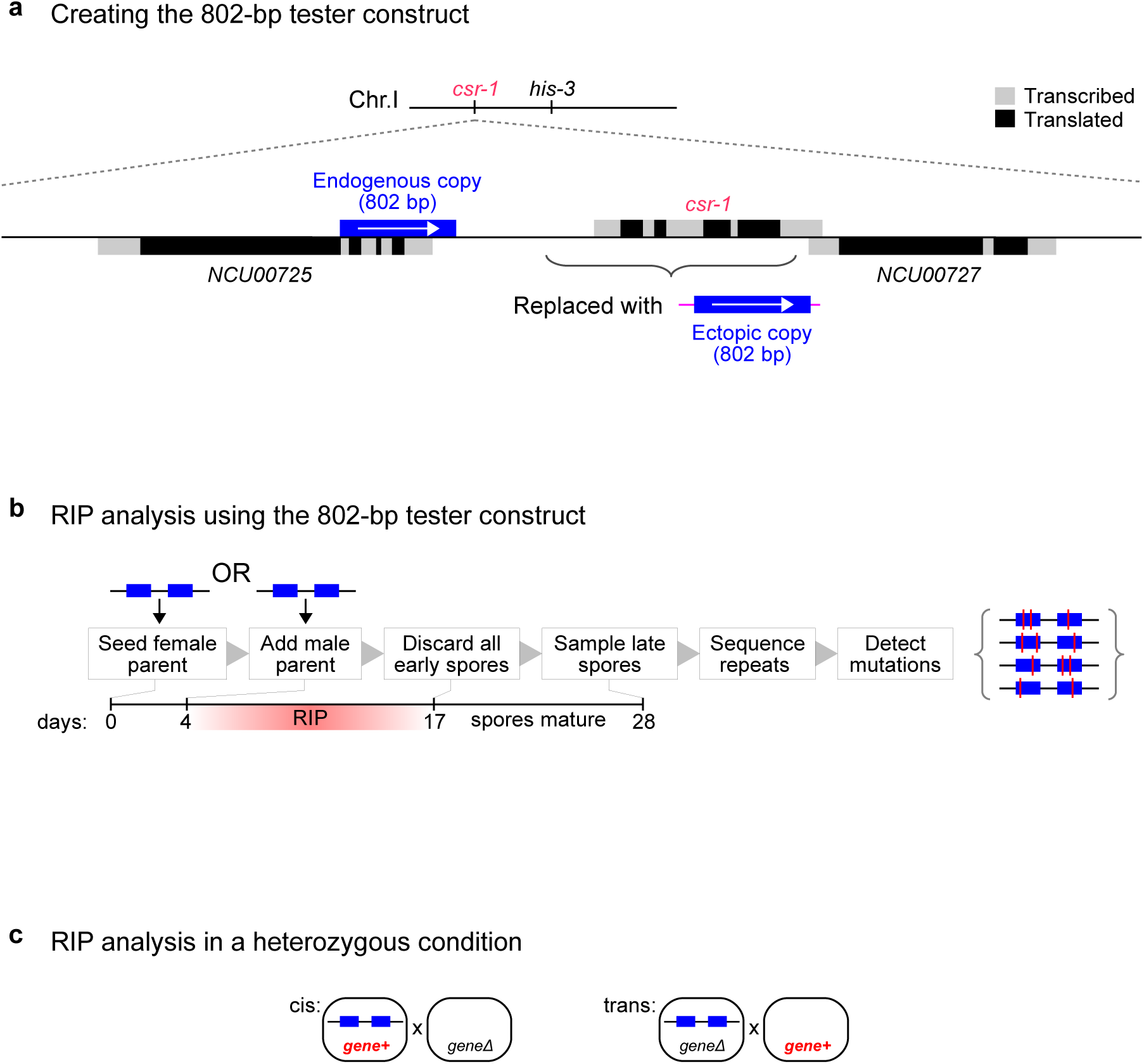
RIP analysis using the 802-bp construct. **a**, The 802-bp tester construct is created by replacing the *csr-1* gene with new DNA containing 802 base-pairs of perfect homology to a neighboring gene *(NCU00725)*. **b**, RIP is detected by sequencing the entire construct in individual “late-arising” spore clones. **c**, When a gene of interest is analyzed in the heterozygous condition for its role in RIP, two configurations are possible: in cis, the wildtype allele is provided together with the tester construct in the same parental nucleus (which could be maternal or paternal); in trans, the wildtype allele is provided by one parent, whereas the tester construct is provided by the other parent (along with the corresponding deletion allele).

**Supplementary Information Figure 2.**
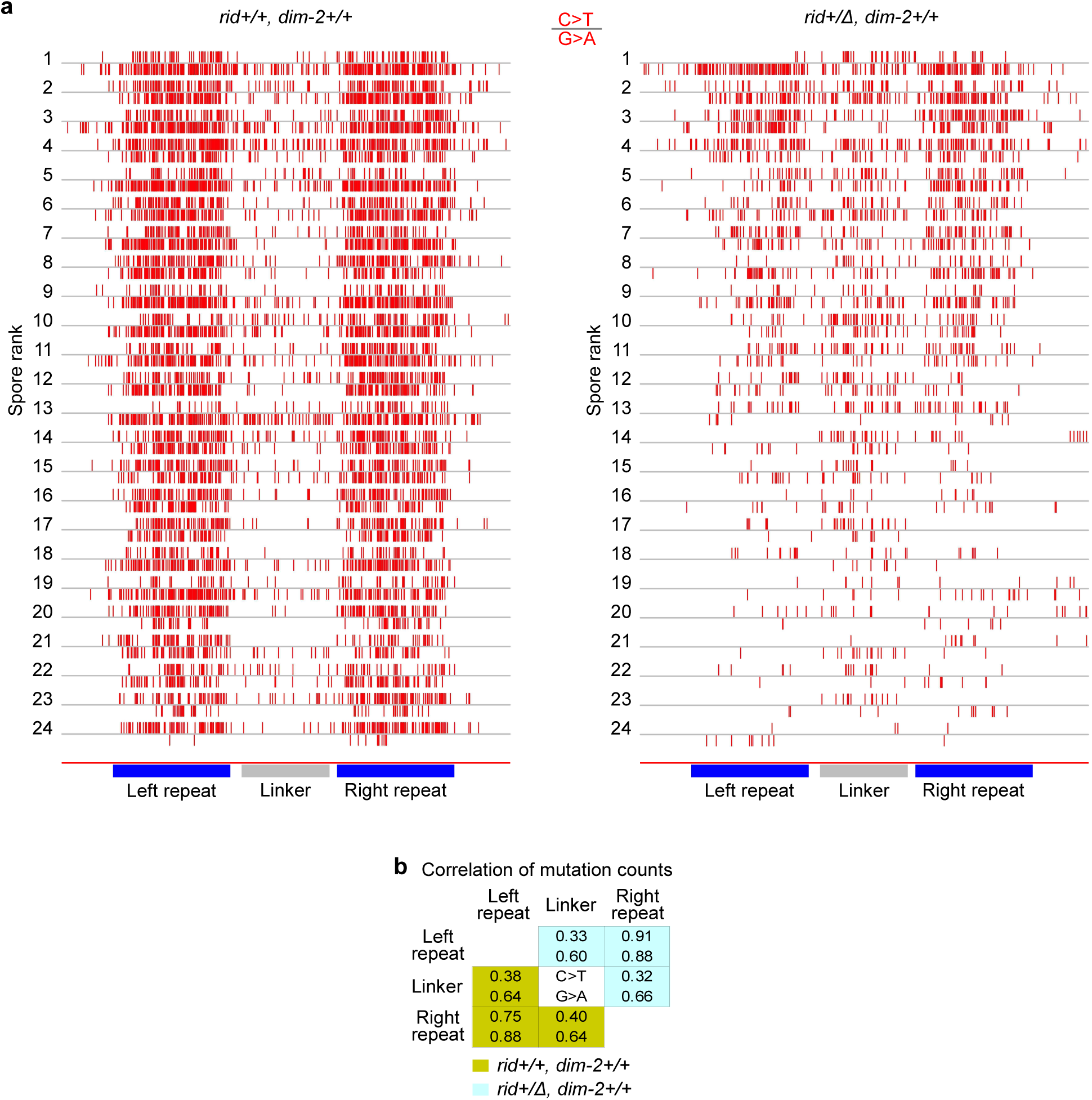
Pair-wise correlations of mutation counts in the linker region and the repeat copies. **a**, All individual C-to-T (“C>T”, “upward” tick-marks) and G-to-A (“G>A”, “downward” tick-marks) mutations are identified in the sample of 24 spores. Spores are ranked by the total number of mutations (C-to-T + G-to-A). Left: cross X2, right: cross X6. **b**, Pearson product-moment correlation coefficients, calculated independently for C-to-T (“C>T”) and G-to-A (“G>A”) mutations, for both crosses in **a**. Only mutations in three regions of interest (802-bp repeats and the 600-bp segment of the linker) were analyzed.

**Supplementary Information Figure 3.**
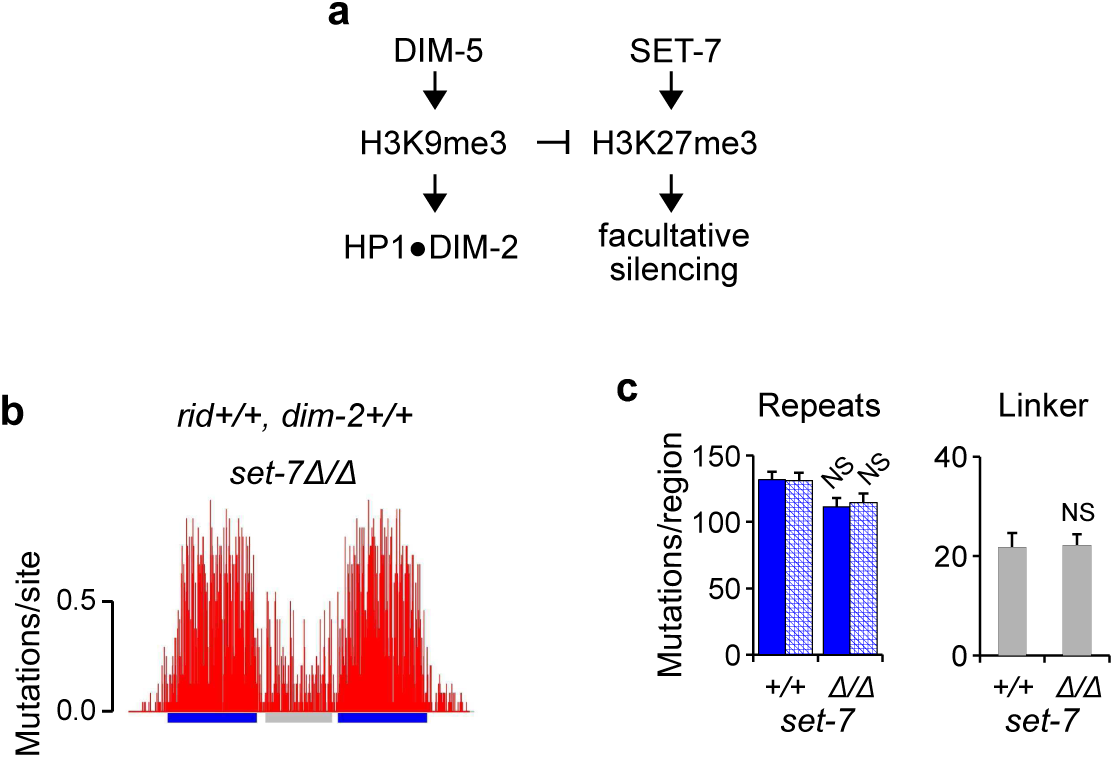
H3K27 methylase SET-7 does not play a detectable role in RIP. **a**, DIM-5-mediated H3K9me3 restricts spreading of SET-7-mediated H3K27me3 in *N. crassa*^22,23^. The lack of SET-7 partially rescues the inability of *dim*-5Δ strains to enter the sexual stage. **b**, RIP mutation profile of the 802-bp construct in the *set*-7Δ/Δ background. Cross X28[N=24]. **c**, The mean number of RIP mutations for the *set*-7Δ/Δ cross in **b** (analyzed as in Fig. 1d), as compared to the reference wildtype cross (X2, Fig. 1d).

**Supplementary Information Figure 4.**
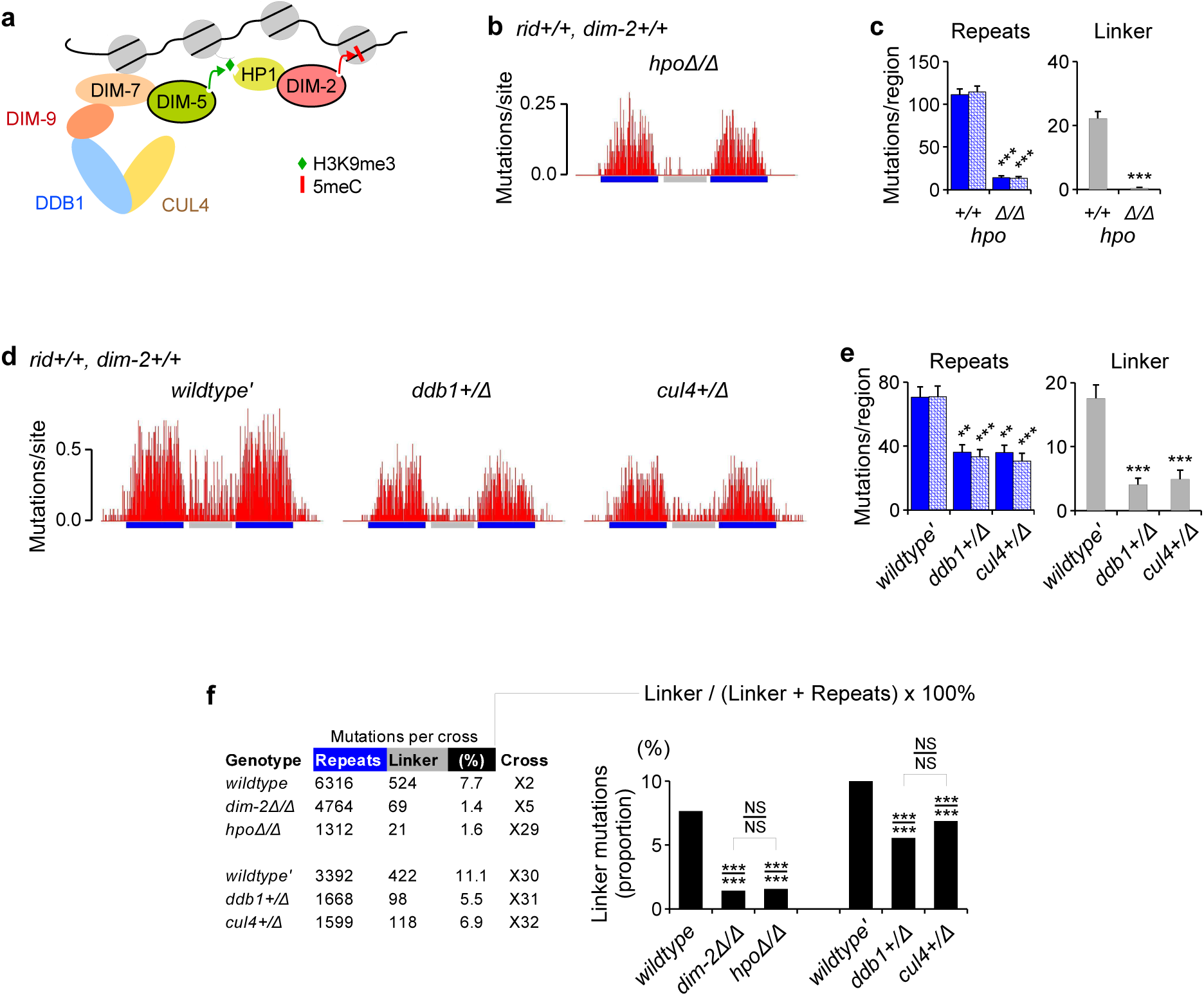
Components of the heterochromatin pathway are involved in DIM-2-mediated RIP. **a**, A cartoon representation of the canonical heterochromatin pathway in *N. crassa:* H3K9me3 is catalyzed by the SUV39 methylase DIM-5 in a complex with DIM-7, DIM-9, DDB1 and CUL4 (ref. 7). Heterochromatin Protein 1 (HP1) recognizes H3K9me3 and directly recruits DIM-2 to DNA. **b**, RIP mutation profile of the 802-bp construct in the *hpo*Δ/Δ cross. Cross X29[N=48]. **c**, The mean number of RIP mutations for the *hpo*Δ/Δ cross in **b** (analyzed as in Fig. 1d), compared to a reference *hpo+/+* cross X28 (Supplementary Information Figure 3). **d**, RIP mutation profiles of the 802-bp construct. From left to right: crosses X30[N=24], X31[N=24], X32[N=24]. **e**, The mean number of RIP mutations for the crosses in **d** (analyzed as in Fig. 1d). **f**, The proportion of “linker” mutations is used as a relative measure of DIM-2-mediated RIP. Proportions are expressed as percentages and compared for significance as raw counts using the chi-squared homogeneity test (*P*-value is indicated above the line) and the Fisher's exact test (*P*-value is indicated below the line). Error bars represent standard error (s.e.m.). Distributions of per-spore mutation counts are compared separately for each designated region by the Kolmogorov-Smirnov test. *** P ≤ 0.001, ** 0.001 < P ≤ 0.01, * 0.01 < P ≤ 0.05, NS 0.05 < P.

**Supplementary Information Figure 5.**
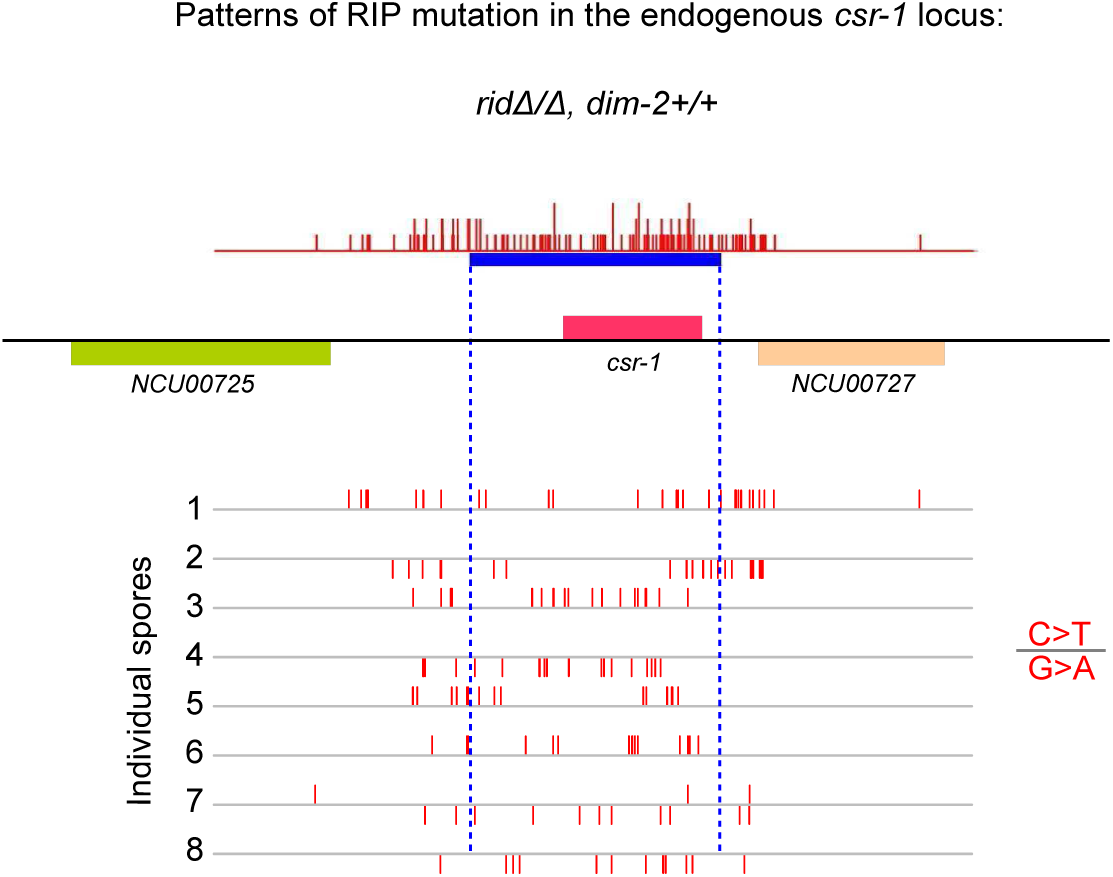
DIM-2-mediated RIP mutations spread towards nearby genes. Eight progeny spores with especially strong levels of DIM-2-mediated RIP were analyzed further for mutation within a larger region. Positions of neighboring genes (corresponding to their start and the stop cordons) are indicated. The endogenous copy of the *csr-1* gene is shown in blue.

**Supplementary Information Figure 6.**
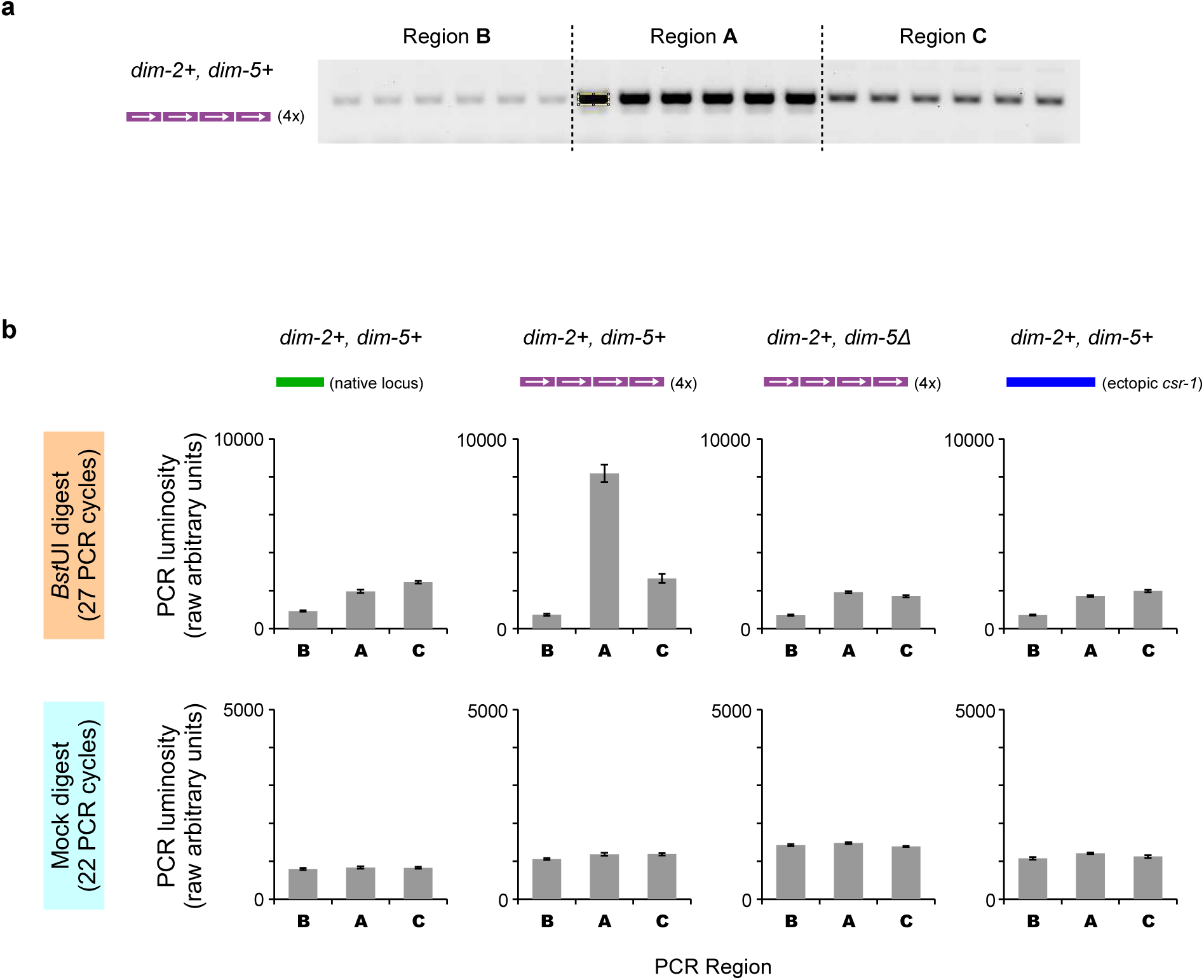
Four tandemly-repeated copies of the dsRed gene induce cytosine methylation in vegetative cells of *N. crassa*. **a**, A gel image with 18 individual PCR products corresponding to the three analyzed regions in Fig. 4a. This particular experiment assays PstUI-digested DNA of the strain T485.4h (Supplementary Information Table 1), which carries the 4x array in the *dim*-2+, *dim*-5+ background. A region of interest (ROI) used to quantify PCR yields is shown in yellow. **b**, Semi-quantitative PCR yields for the three analyzed regions (Fig. 4a) are reported in raw luminosity units (mean ± standard deviation). Top row: DNA was digested with *Bst*UI, bottom row: DNA was mock-digested. Assayed strains (from left to right): FGSC#9720, T485.4h, T486.3h and T402.1h (Supplementary Information Table 1). The presence of intact repetitive DNA was verified by PCR in mock-digested samples using primers *NcHis3_F4* and *NcHis3 R1* (Supplementary Information Table 3).

## MATERIALS AND METHODS

### Plasmids

Plasmids pEAG66, pEAG115A, pEAG186B, pEAG186K and pEAG186L are based on the plasmid pCSR1^31^. Plasmids pEAG82B (inactive copy of the *csr*-1 gene), pEAG236A (the “4x” construct), pEAG244B (active copy of the *dim*-5 gene) and pEAG244G (inactive copy of the *dim*-5 gene) are based on the plasmid pMF280^32^ and pMF334^33^. Annotated maps of pEAG66, pEAG115A, pEAG186B, pEAG186K and pEAG186L were published previously^10^. Annotated maps of pEAG82B, pEAG236A, pEAG244B and pEAG244G are provided in Supplementary Information File 1 (in GenBank format, individual plain-text files are compressed with tar/gz).

### Manipulation of *Neurospora* strains

All strains used in this study are listed in Supplementary Information Table 1. All *FGSC#* strains were obtained from the Fungal Genetics Stock Center^34^. Transformation of the *csr*-1 locus using linearized plasmids pEAG66, pEAG115A, pEAG186B, pEAG186K and pEAG186L was carried out as described previously^10^. Transformation of the *his*-3 locus using linearized plasmids pEAG82B, pEAG236A, pEAG244B and pEAG244G was done analogously, by electroporation of macroconidia. Homokaryotic *his*-3+ strains were isolated from primary heterokaryotic transformants by macroconidiation. Crosses were setup on agar slants in capped culture tubes (15 mm x 125 mm) as described previously^10^. 0.25 mg/ml histidine was added to the crossing medium only if required by the female parent. All crosses analyzed in this study are listed in Supplementary Information Table 2.

### Analysis of the cyclosporin-resistant phenotype

This assay was designed to detect very low levels of RIP activity involving two widely-separated repeats of the *csr*-1 gene^12^. A copy of the *csr*-1 gene (including the promoter region) was integrated into the *his*-3 locus, 2.7 million base-pairs away from the endogenous *csr*-1. The ectopic copy carries a single G-to-A mutation in the 5' splice site, which confers resistance to cyclosporin. Thus, inactivation of the endogenous *csr*-1 allele was sufficient to confer a cyclosporin-resistant phenotype. All ejected spores (both early- and late-arising) were collected into 1 ml of distilled water. Spores were heat-shocked for 30 minutes at 60°C to induce germination. To estimate the total number of viable spores, 10 μl of the original 1-ml suspension were diluted 100-fold (back to 1 ml) and 50 μl of that 1:100 dilution were plated on non-selective sorbose agar solidified in standard Petri dishes (100 mm × 15 mm). 3 replica dishes were plated in total, consuming 3×50 μl of the 1:100 dilution. The number of viable spores was estimated as the average number of colonies per dish multiplied by the overall dilution factor. To estimate the number of cyclosporin-resistant progeny in crosses with strong RIP, additional 3×50 μl From the same 1:100 dilution were plated in triplicate on sorbose agar containing cyclosporin A (5 μg/ml; Sigma cat. no. 30024-25MG). The total number of cyclosporin-resistant progeny was estimated by the same formula. In cases of weak RIP (expected in the absence of RID or in the absence of the ectopic *csr*-1 copy), the remaining 0.99 ml of the original 1:1 suspension were plated directly on selective medium, and the number of cyclosporin-resistant progeny was determined by the total count of growing colonies.

### Sampling RIP products for sequence analysis

RIP is likely activated in haploid parental nuclei that become assorted into dikaryotic ascogenous cells^35^. Approximately 6-7 rounds of RIP may occur as these pre-meiotic nuclei continue to divide by mitosis. After the nuclei undergo karyogamy and meiosis, the haploid meiotic products are packaged into progeny spores and ejected from the mature perithecium (the fruiting body). RIP mutation tends to be intrinsically weak in the “early-arising” spores (ejected within the first 1-3 days after the onset of sporogenesis) but then increases in “late-arising” spores (ejected over the next 7-10 days). 100-200 spore-producing perithecia can normally develop in a single test tube under our experimental conditions^10^. Each perithecium represents an autonomous anatomical structure; it hosts several independent lineages of ascogenous cells and ultimately produces several hundred progeny spores. All ejected spores correspond to one statistical population. Late-arising spores are sampled without any knowledge of their RIP status. Because the number of sampled spores (24-60 from each cross) is much smaller than the estimated number of ascogenous lineages, each spore effectively represents an independent measure of RIP activity. Despite the apparent spore-to-spore variability in RIP levels, our studies has indicated that the expected number of RIP mutations (the arithmetic mean) represents an accurate and useful measure of RIP activity^10,11^.

### Genomic DNA extraction, PCR amplification, and sequencing

Genomic DNA was extracted from individual spore clones as described previously^10^. PCR products were purified using Omega Bio-tek kit and sequenced directly by primer walking. The following primers were used for PCR and sequencing (primers are provided in Supplementary Information Table 3): The 802-bp construct was amplified with primers *P66_Seq3* and *RIP2 _R1;* sequenced with primers *P66_Seq1, P66_Seq3*, *LNK SeqRl* and *RIP2 R2.* All (220+200)-bp constructs were amplified with primers *P66_Seq3* and *RIP2_R1;* sequenced with primers *P66_Seq17* and *P66_Seq18.* The entire 4x construct was amplified with primers *NcHis3_R6* and *NcHis3_F7;* sequenced with primers *NcHis3_F4, NcHis3_F7, NcHis3_R1, NcHis3_R4*, *NcHis3_R6, P236A_Seq1, P236A_Seq2, P236A_Seq4, P236A_Seq5* and *P236A_Seq8.* If the entire 4x construct could not be amplified as a single fragment, piece-wise applications were carried out using pair-wise combinations of the above sequencing primers. The endogenous *csr-1* gene was amplified with primers *P66_Seq10* and *RIP2_R1;* sequenced with primers *P66_Seq10, P66_Seq12, CSRI _SeqF, CSRI_SeqR* and *RIP2_R1*. Eight progeny spores with particularly strong levels of DIM-2-mediated mutation were selected for further sequence analysis of the adjoining genomic regions. For these eight clones, the extended 5.4-kbp region containing the endogenous *csr-1* gene was amplified with primers *P66_Seq3* and *RIP2_R3;* and sequenced with additional primers *P66_Seq3, P66_Seq9, P66_Seq4, CSR1_SeqR2, CSR1_SeqR3* and *RIP2_R3.* If a primer site appeared affected by RIP, additional *ad hoc* primers were used instead. Sequencing reactions were read with an ABI3730xl DNA analyzer at the DNA Resource Core of Dana-Farber/Harvard Cancer Center (funded in part by NCI Cancer Center support grant 2P30CA006516-48). Individual chromatograms were assembled into contigs with Phred/Phrap. All assembled contigs were validated manually using Consed.

### Sequence and statistical analysis

For each cross, assembled repeat-containing contigs were aligned with the parental (reference) sequence using ClustalW (all alignments analyzed in this study are provided in Supplementary Information File 2 in ClustalW format, individual plain-text files are compressed with tar/gz). Mutations were detected and analyzed as described previously^10^. The absolute level of RIP in a given region of interest was expressed (1) as the expected number of mutations (the arithmetic mean) and, if necessary, (2) as the percentage of mutated strands, where all C-to-T mutations were considered to be on the top strand and all G-to-A mutations - on the bottom strand^10^. Standard error of the mean (s.e.m.) was used as a measure of variation. The following regions were analyzed for the 802-bp construct: the endogenous “left” repeat copy (positions 348-1149), the 600-bp segment of the linker (positions 1227-1826), and the ectopic “right” repeat copy (positions 1879-2680). The following region was analyzed for the (220+200)-bp construct: the 450-bp segment of the linker region (positions 622-1071). Empirical distributions of mutation counts (i.e., the numbers of C-to-T and G-to-A mutations found together within a given region on per-spore basis) were compared by the Kolmogorov-Smirnov test^10^. The pair-wise differences in the percentage of mutated strands and the proportion of linker mutations were compared for significance as the original raw counts by (1) Pearson's Chi-squared homogeneity test with Yates' continuity correction and (2) Fisher's exact test (as implemented in R).

### Analysis of cytosine methylation in vegetative cells of *N. crassa*

#### Purification of genomic DNA

Crude preparations of genomic DNA were obtained by the method used for extracting DNA for RIP analysis (above; ref. 10), except that phenol/chloroform was replaced with chloroform in all purification steps. DNA was precipitated with isopropanol, washed with ethanol, resuspended in 80 μ1 of water and mixed with 20 μ1 of the 5x loading buffer (20% v/v glycerol, 0.25% SDS, 5 mM EDTA, 1 mg/ml xylene cyanol). The entire sample (100 μ1 was loaded into a single well of a 0.75% agarose mini-gel (NuSieve GTG Agarose, cat. no. 50080) containing ethidium bromide. The gel was run at 3 V/cm in 1× TAE (Tris-Acetate-EDTA) buffer for 1.5 hours. A region of the gel containing high-molecular weight DNA was excised with a sterile razor blade under long UV light and digested with β-Agarase I (NEB, cat. no. M0392L; 3 hours at 42°C). Following a single extraction with chloroform, genomic DNA was precipitated with ammonium acetate/ethanol, washed with ethanol, and resuspended in 50 μl of water. DNA concentrations were measured using NanoDrop 2000. A typical procedure yielded 0.5-1.0 μg of purified DNA (per one mycelial sample grown overnight in 2 ml of 1x Vogel's culture medium).

#### Restriction digest with BstUI

*Bst*UI recognition sequence (5'-CGCG-3') contains four cytosines that can be potentially methylated by DIM-2 (two cytosines on the top strand, and two cytosines on the bottom strand); and methylation of any one (or more) of these cytosines protects the site from cleavage by *Bst*UI^36^. Each restriction reaction was setup in the total volume of 50 μ1, using 0.2 μg of purified genomic DNA from the previous step and 4 μl of *Bst*UI (10 U/μ1 NEB, cat. no. R0518S). Reactions were incubated for 3 hours at 50°C. *Bst*UI was inactivated by Proteinase K (by adding 2 μl of Proteinase solution, 20 mg/ml, and incubating at 50°C for additional 1.5 hours). Proteinase K was heat-inactivated (95°C for 10 min) and the volume of each sample was adjusted to 200 μl (by adding 148 μl of water). Mock-digested samples were processed in exactly the same way, except that 4 μl of 50% glycerol were added to each reaction instead of *Bst*UI.

#### Semi-quantitative PCR

Each PCR was run in the total volume of 10 μ1 Three genomic regions were amplified separately for each DNA sample: Region A (primers *REG_A_F1* and *REG_A_R1)*, Region B (primers *REG_B_F1* and *REG_B_R1)* and Region C (primers *REG_C_F1* and *REG_C_R1).* DNA sample concentrations were adjusted (by no more than a factor of 1.5) to produce comparable PCR yields of Region B. No additional adjustments were implemented. Each DNA sample (corresponding to 20-36 μl of the 200μl product from the previous step) was used to make one Principal Master Mix (PMM) for 18 individual PCRs. Each PMM also received 18 μl of the 10x reaction buffer (EconoTaq DNA Polymerase, Lucigen, cat. no. F93366-1), 7.2 μl of the dNTP solution (NEB, cat. no. N0447S), and 5.4 μl of EconoTaq polymerase (in the total volume of 162 μ1. Each PMM was split equally into three 54-μl Region-Specific Master Mixes (RSMMs). Each RSMM received 6 μl of a primer solution (containing one pair of primers corresponding to a given PCR region, at 5 pmol/μl each), bringing its total volume to 60 μ1 Each RSMM was then split into six individual 10-μl reactions. All PCRs were run in a C1000 Touch thermal cycler (Bio-Rad) using the following program: 94°C-2', {94°C-20"; 59°C-20''; 72°C-20''} × N, 72°C-2'. For all digested DNA samples, the value of N (the number of amplification cycles) was set to 27. For all mock-digested samples, the value of N was set to 22. The number of cycles was determined empirically to ensure that all PCRs remained in the exponential regime. Each PCR product was mixed with 2 μ1 of the 5x buffer (above) and loaded on a 2% regular agarose gel in the total volume of 12 μ1. All PCRs corresponding to a given DNA sample were analyzed side-by-side on the same gel. Gels were run for 30 min in 0.5x TAE at 6 V/cm and scanned with the resolution of 50 microns using Molecular Imager FX (Bio-Rad). Luminosities of PCR products were measured in ImageJ (with “Analyze → Measure”). Same region-of-interest (ROI) parameters were used for quantifying all PCR bands within one image. A constant value corresponding to the image background (estimated separately based on five independent measurements) was subtracted from a raw luminosity value of each PCR band. The relative number of uncleavable *Bst*UI sites in the digested DNA sample (in Region A) was estimated using a calibration curve based on a series of two-fold dilutions of the corresponding mock-digested DNA sample.

---

**Supplementary Information Table 1.** *N. crassa* strains used in this study. All inserts were validated by PCR and sequencing; all gene deletions were validated by diagnostic PCR. Primer sequences are provided in Supplementary Information Table 3. Tester repeats are shown in bold.

| Strain | Mating type | Genotype | Source | Reference |
| --- | --- | --- | --- | --- |
| TGSC#2489 | A | wildtype | The Fungal Genetics Stock Center | ref. 34 |
| FGSC#4200 | a | wildtype | The Fungal Genetics Stock Center | ref. 34 |
| FGSC#8594 | a | *dim-2Δ, his-3* | The Fungal Genetics Stock Center | ref. 34 |
| FGSC#9539 | a | *mus-52Δ, his-3* | The Fungal Genetics Stock Center | ref. 34 |
| FGSC#9720 | A | *mus-52Δ, his-3* | The Fungal Genetics Stock Center | ref. 34 |
| FGSC#11198 | a | *ddb1Δ* | The Fungal Genetics Stock Center | ref. 34 |
| FGSC#12374 | a | *cul4Δ* | The Fungal Genetics Stock Center | ref. 34 |
| FGSC#12353 | a | *ridΔ* | The Fungal Genetics Stock Center | ref. 34 |
| FGSC#12354 | A | *ridΔ* | The Fungal Genetics Stock Center | ref. 34 |
| FGSC#12440 | A | *spo11Δ* | The Fungal Genetics Stock Center | ref. 34 |
| FGSC#14522 | a | *hpoΔ* | The Fungal Genetics Stock Center | ref. 34 |
| FGSC#15885 | a | *dim-5Δ* | The Fungal Genetics Stock Center | ref. 34 |
| S-353 | a | *dim-5Δ, set-7Δ* | Zachary Lewis, University of Georgia, USA | ref. 23 |
| S-354 | A | *dim-5Δ, set-7Δ* | Zachary Lewis, University of Georgia, USA | ref. 23 |
| T103.3 | A | **802-bp**, *mus-52Δ, his-3* | FGSC#9720 transformed with pEAG115A | ref. 10 |
| T208.3 | A | **RANDOM**, *mus-52Δ, his-3* | FGSC#9720 transformed with pEAG186B | ref. 10 |
| T221.18 | A | **4H-7N**, *mus-52Δ, his-3* | FGSC#9720 transformed with pEAG186K | ref. 10 |
| T223.4 | A | **3H-8N**, *mus-52Δ, his-3* | FGSC#9720 transformed with pEAG186L | ref. 10 |
| T337.1h | a | **csr-1(Dp)::his-3+**, *ridΔ, mus-52Δ* | C03.1 transformed with pEAG82B | this study |
| T364.6 | A | **802-bp**, *ridΔ* | FGSC#12354 transformed with pEAG115A | this study |
| T382.32 | a | **802-bp**, *dim-2Δ, his-3* | FGSC#8594 transformed with pEAG115A | this study |
| T402.1h | A | **csr-1(Dp)::his-3+**, *mus-52Δ* | FGSC#9720 transformed with pEAG82B | this study |
| T451.6h | A | **4x::his-3+**, *ridΔ, mus-52Δ* | C02.1 transformed with pEAG236A | this study |
| T452.1h | A | **4x::his-3+**, *ridΔ, dim-2Δ, mus-52Δ* | C58.40 transformed with pEAG236A | this study |
| T475.6h | a | *ridΔ, dim-5Δ, set-7Δ, mus-52Δ, his-3::dim-5+* | C146.1 transformed with pEAG244B | this study |
| T482.3 | A | **RANDOM**, *ridΔ, mus-52Δ, his-3* | C02.1 transformed with pEAG186B | this study |
| T484.1 | A | **3H-8N**, *ridΔ, mus-52Δ, his-3* | C02.1 transformed with pEAG186L | this study |
| T485.4h | A | **4x::his-3+**, *mus-52Δ* | FGSC#9720 transformed with pEAG236A | this study |
| T486.3h | a | **4x::his-3+**, *ridΔ, dim-5Δ, set-7Δ, mus-52Δ* | C146.1 transformed with pEAG236A | this study |
| T487.3 | A | **4H-7N**, *ridΔ, mus-52Δ, his-3* | C02.1 transformed with pEAG186K | this study |
| T488.1h | a | *ridΔ, dim-5Δ, set-7Δ, mus-52Δ, his-3::dim-5(Y283F)* | C146.1 transformed with pEAG244G | this study |
| T489.1 | A | **NO-REPEAT**, *mus-52Δ, his-3* | FGSC#9720 transformed with pEAG66 | this study |
| C02.1 | A | *ridΔ, mus-52Δ, his-3* | Cross progeny of FGSC#9539 and FGSC#12354 | this study |
| C03.1 | a | *ridΔ, mus-52Δ, his-3* | Cross progeny of FGSC#9720 and FGSC#12353 | this study |
| C50.3 | A | *ridΔ, dim-2Δ* | Cross progeny of FGSC#8594 and FGSC#12354 | this study |
| C52.1 | a | **csr-1(Dp)::his-3+**, *ridΔ, dim-2Δ* | Cross progeny of C50.3 and T337.1h | this study |
| C57.8 | A | **802-bp**, *ridΔ, dim-2Δ* | Cross progeny of C52.1 and T364.6 | this study |
| C58.7 | a | *ridΔ, dim-2Δ, mus-52Δ, his-3* | Cross progeny of C50.3 and C03.1 | this study |
| C58.40 | A | *ridΔ, dim-2Δ, mus-52Δ, his-3* | Cross progeny of C50.3 and C03.1 | this study |
| C96.1 | A | *dim-2Δ* | Cross progeny of FGSC#8594 and FGSC#2489 | this study |
| C97.2 | A | **802-bp**, *ridΔ, hpoΔ* | Cross progeny of FGSC#14522 and T364.6 | this study |
| C97.6 | a | **802-bp**, *hpoΔ* | Cross progeny of FGSC#14522 and T364.6 | this study |
| C98.18 | A | **802-bp**, *ridΔ, dim-5Δ* | Cross progeny of FGSC#15885 and T364.6 | this study |
| C135.3 | A | **802-bp**, *set-7Δ, spo11Δ, hpoΔ* | Cross progeny of XG1.2 and FGSC#12440 | this study |
| C139.10 | a | *ridΔ, set-7Δ, mus-52Δ, his-3* | Cross progeny of C58.7 and S-354 | this study |
| C146.1 | a | *ridΔ, dim-5Δ, set-7Δ, mus-52Δ, his-3* | Cross progeny of XG3.4 and C139.10 | this study |
| C146.3 | a | *ridΔ, dim-5Δ, set-7Δ* | Cross progeny of XG3.4 and C139.10 | this study |
| R88.6 | A | **csr-1(Dp)::his-3+** | Cross progeny of C52.1 and FGSC#2489 | this study |
| XG1.2 | a | **802-bp**, *set-7Δ, dim-5Δ, hpoΔ* | Cross progeny of C97.2 and S-353 | this study |
| XG1B.17 | a | *set-7Δ* | Cross progeny of C97.2 and S-353 | this study |
| XG3.4 | A | **802-bp**, *dim-5Δ, set-7Δ* | Cross progeny of C97.6 and S-354 | this study |
| XG3.7 | A | **802-bp**, *set-7Δ* | Cross progeny of C97.6 and S-354 | this study |
| XG3B.11 | a | *set-7Δ, hpoΔ* | Cross progeny of C97.6 and S-354 | this study |
| XG26.8 | A | **802-bp**, *dim-5Δ, set-7Δ, ridΔ* | Cross progeny of C98.18 and C146.3 | this study |

**Supplementary Information Table 2.** Crosses analyzed in this study. **Figure**, Figure/panel with a corresponding RIP mutation profile; **Cross**, a unique cross identifier used in this study; **Cross genotype**, provided as pairs of maternal/paternal alleles; **Parental strains**, strain genotypes are provided in Supplementary Information Table 1. Strains with the assayed repeat constructs are shown in bold; **N(S)**, the total number of progeny spores analyzed (by sequencing) for a given cross/condition. All sequence alignments are provided in Supplementary Information File 2. Sequence data for crosses X2, X20, X21 and X22 were published previously (ref. 10) and are provided in File 2 only for the sake of convenience.

|  |  | Cross genotype |  | Parental strains |  |  |  |
| --- | --- | --- | --- | --- | --- | --- | --- |
| Figure | Cross | *rid, dim-2* | *other* | Female | Male | N(S) | Reference |
| Fig. 1 |  |  |  |  |  |  |  |
| c | X1 | *rid*+/*+*, *dim-2*+/*+* |  | FGSC#4200 | **T489.1** | 48 | this study |
| d | X2 | *rid*+/*+*, *dim-2*+/*+* |  | FGSC#4200 | **T103.3** | 24 | ref. 10 |
| d | X3 | *rid*Δ/Δ, *dim-2*+/*+* |  | T337.1h | **T364.6** | 48 | this study |
| d | X4 | *rid*Δ/Δ, *dim-2*Δ/Δ |  | C52.1 | **C57.8** | 48 | this study |
| d | X5 | *rid*+/*+*, *dim-2*Δ/Δ |  | C96.1 | **T382.32** | 24 | this study |
| d | X6 | *rid*+/*Δ*, *dim-2*+/*+* |  | FGSC#4200 | **T364.6** | 24 | this study |
| d | X7 | *rid*+/*+*, *dim-2*+/*Δ* |  | FGSC#2489 | **T382.32** | 24 | this study |
| Fig. 2 |  |  |  |  |  |  |  |
| a | X8 | *rid*Δ/Δ, *dim-2*+/*+* | *dim-5*Δ/*+*, *set-7*Δ/*+* | C146.3 | **T364.6** | 60 | this study |
| a | X9 | *rid*Δ/Δ, *dim-2*+/*+* | *dim-5*Δ/Δ, *set-7*Δ/*+* | C146.3 | **C98.18** | 60 | this study |
| b | X10 | *rid*Δ/Δ, *dim-2*+/*+* | *dim-5*Δ/Δ, *set-7*Δ/Δ | **XG26.8** | C146.1 | 60 | this study |
| b | X11 | *rid*Δ/Δ, *dim-2*+/*+* | *dim-5*Δ/*+*, *set-7*Δ/Δ | **XG26.8** | C139.10 | 60 | this study |
| b | X12 | *rid*Δ/Δ, *dim-2*+/*+* | *his-3::dim-5*+, *dim-5*Δ/Δ, *set-7*Δ/Δ | **XG26.8** | T475.6h | 60 | this study |
| b | X13 | *rid*Δ/Δ, *dim-2*+/*+* | *his-3::dim-5* (Y283F), *dim-5*Δ/Δ, *set-7*Δ/Δ | **XG26.8** | T488.1h | 60 | this study |
| d | X14 | *rid*+/*Δ*, *dim-2*+/*+* | *dim-5*+/*+*, *set-7*Δ/*+* | XG1B.17 | **T364.6** | 24 | this study |
| d | X15 | *rid*+/*Δ*, *dim-2*+/*+* | *dim-5*Δ/*+*, *set-7*Δ/*+* | S-353 | **T364.6** | 24 | this study |
| d | X16 | *rid*+/*Δ*, *dim-2*+/*+* | *dim-5*Δ/Δ, *set-7*Δ/*+* | S-353 | **C98.18** | 48 | this study |
| Fig. 3 |  |  |  |  |  |  |  |
| d | X17 | *rid*Δ/Δ, *dim-2*+/*+* |  | T337.1h | **T482.3** | 60 | this study |
| d | X18 | *rid*Δ/Δ, *dim-2*+/*+* |  | T337.1h | **T484.1** | 60 | this study |
| d | X19 | *rid*Δ/Δ, *dim-2*+/*+* |  | T337.1h | **T487.3** | 60 | this study |
| d | X20 | *rid*+/*+*, *dim-2*+/*+* |  | FGSC#4200 | **T208.3** | 60 | ref. 10 |
| d | X21 | *rid*+/*+*, *dim-2*+/*+* |  | FGSC#4200 | **T223.4** | 90 | ref. 10 |
| d | X22 | *rid*+/*+*, *dim-2*+/*+* |  | FGSC#4200 | **T221.18** | 60 | ref. 10 |
| Fig. 4 |  |  |  |  |  |  |  |
| d | X23 | *rid*+/*+*, *dim-2*+/*+* |  | **R88.6** | FGSC#4200 | 10 | this study |
| d | X24 | *rid*Δ/Δ, *dim-2*+/*+* |  | **T337.1h** | C02.1 | 70 | this study |
| d | X25 | *rid*Δ/Δ, *dim-2*Δ/Δ |  | C50.3 | **C52.1** | 28 | this study |
| Fig. 5 |  |  |  |  |  |  |  |
| b | X26 | *rid*+/*Δ*, *dim-2*Δ/Δ |  | FGSC#8594 | **T452.1h** | 14 | this study |
| b | X27 | *rid*Δ/Δ, *dim-2*+/*+* |  | C03.1 | **T451.6h** | 30 | this study |
| SI Fig. 3 |  |  |  |  |  |  |  |
| b | X28 | *rid*+/*+*, *dim-2*+/*+* | *set-7*Δ/Δ | XG1B.17 | **XG3.7** | 24 | this study |
| SI Fig. 4 |  |  |  |  |  |  |  |
| b | X29 | *rid*+/*+*, *dim-2*+/*+* | *hpo*Δ/Δ, *set-7*Δ/Δ, *spo11*Δ/*+* | **C135.3** | XG3B.11 | 48 | this study |
| d | X30 | *rid*+/*+*, *dim-2*+/*+* | *set-7*Δ/*+* | **XG3.7** | FGSC#4200 | 24 | this study |
| d | X31 | *rid*+/*+*, *dim-2*+/*+* | *ddb1*+/*Δ*, *set-7*Δ/*+* | **XG3.7** | FGSC#11198 | 24 | this study |
| d | X32 | *rid*+/*+*, *dim-2*+/*+* | *cul4*+/*Δ*, *set-7*Δ/*+* | **XG3.7** | FGSC#12374 | 24 | this study |

**Supplementary Information Table 3.**
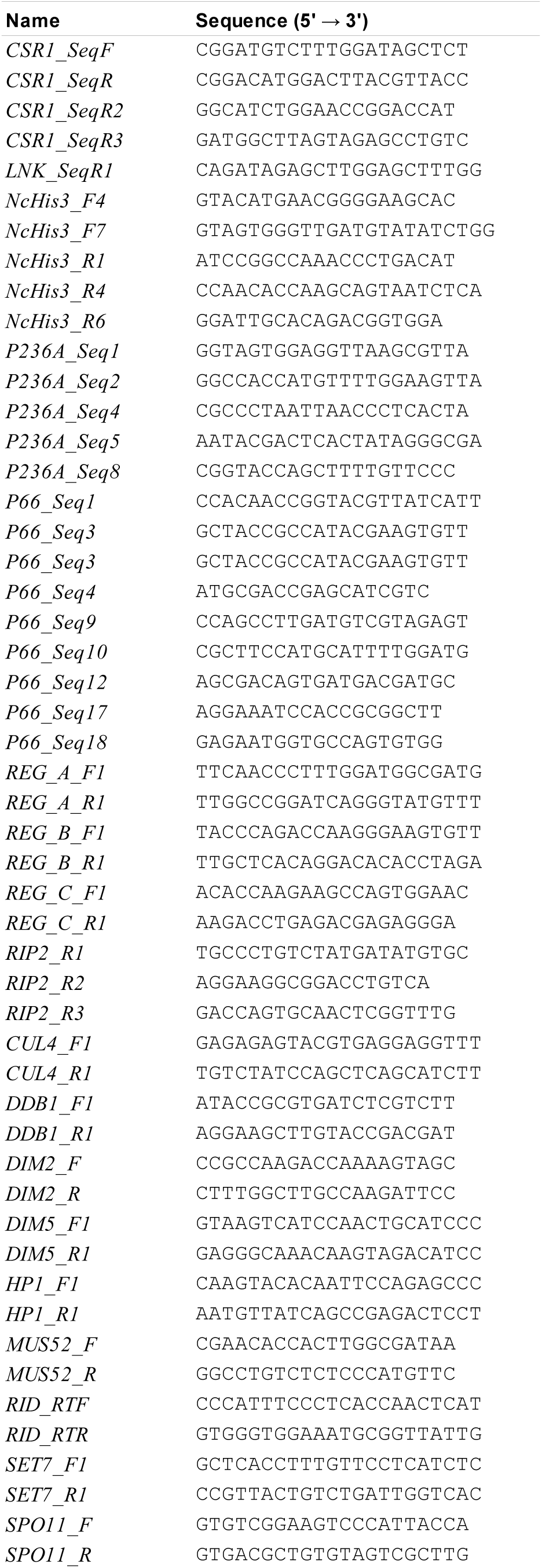
Primers used in this study

